# Decoil: Reconstructing extrachromosomal DNA structural heterogeneity from long-read sequencing data

**DOI:** 10.1101/2023.11.15.567169

**Authors:** Mădălina Giurgiu, Nadine Wittstruck, Elias Rodriguez-Fos, Rocío Chamorro González, Lotte Brückner, Annabell Krienelke-Szymansky, Konstantin Helmsauer, Anne Hartebrodt, Philipp Euskirchen, Richard P. Koche, Kerstin Haase, Knut Reinert, Anton G. Henssen

## Abstract

Circular extrachromosomal DNA (ecDNA) is a form of oncogene amplification found across cancer types and associated with poor outcome in patients. EcDNA can be structurally complex and contain rearranged DNA sequences derived from multiple chromosome locations. As the structure of ecDNA can impact oncogene regulation and may indicate mechanisms of its formation, disentangling it at high resolution from sequencing data is essential. Even though methods have been developed to identify and reconstruct ecDNA in cancer genome sequencing, it remains challenging to resolve complex ecDNA structures, in particular amplicons with shared genomic footprints. We here introduce Decoil, a computational method which combines a breakpoint-graph approach with *LASSO* regression to reconstruct complex ecDNA and deconvolve co-occurring ecDNA elements with overlapping genomic footprints from long-read nanopore sequencing. Decoil outperforms *de-novo* assembly and alignment-based methods in simulated longread sequencing data for both simple and complex ecDNAs. Applying Decoil on whole genome sequencing data uncovered different ecDNA topologies and explored ecDNA structure heterogeneity in neuroblastoma tumors and cell lines, indicating that this method may improve ecDNA structural analyzes in cancer.

## 1 Introduction

Circular extrachromosomal DNA (ecDNA) is an important form of oncogene amplification in cancer [1], which can be formed through multiple mechanisms [2–4] and have a large size (up to several MB [5]). As a result, ecDNA can be structurally diverse, with different functional outcomes. The structure of ecDNA can impact gene regulation through the rearrangement of regulatory elements as well as topologically associated domain (TAD) boundaries [6]. To explore ecDNA diversity and complexity, high-resolution computational methods to reconstruct ecDNA with high accuracy from genome sequencing data are required. The reconstruction of ecDNA from sequencing data remains challenging due to the variable complexity and intratumor heterogeneity of these circular elements. On the one hand, a single ecDNA can be heavily rearranged and contain low-complexity sequence regions (e.g. repeats), which pose a challenge to mapping and *de-novo* assembly based methods. On the other hand, one tumor can contain different ecDNA elements [7, 8], which can either originate from different or shared genomic locations [9]. The latter scenario may be very challenging for ecDNA reconstruction, as different co-occurring ecDNA elements have overlapping genomic footprints, making it difficult to attribute the overlapping features to each of the different circular elements. In the past years, several computational tools have been developed to reconstruct ecDNA from different input data. Some methods were developed to detect circularized DNA regions by identifying the breakpoints leading to circularization (circle-enrich-filter [10], Circle-Map [11], ecc finder [12]). These approaches are suitable for detecting simple circular amplicons, but overlook complex ecDNA structures. To overcome these limitations, more recently, methods focused on reconstructing complex ecDNA based on different technologies, e.g. short-read whole-genome sequencing (AmpliconArchitect [13]), optical-mapping combined with short-read sequencing (AmpliconReconstructor [14]), and long-read sequencing were developed (CReSIL [15]). Lastly, methods have been developed to delineate ecDNA structural heterogeneity [7], by isolating and reconstructing individual ecDNA elements, leveraging *a priori* knowledge about the ecDNA present in the sample of interest. However, a method that reconstructs complex ecDNA structures and captures heterogeneity by distinguishing between ecDNA elements with overlapping genomic footprints from whole-genome sequencing (WGS) data without such *a priori* knowledge is still largely missing to date. We here present Decoil, a computational method to reconstruct genome-wide complex ecDNA elements and deconvolve individiual ecDNAs with shared genomic sequences from bulk whole-genome long-read sequencing using Nanopore technology. Decoil is a graph-based approach integrating the structural variant (SV) and coverage profiles to deconvolve and reconstruct complex ecDNAs. It uses *LASSO* regression to infer likely ecDNA structures and estimate their relative proportions, by accounting for circular elements with overlapping genomic footprints. The model can separate individual ecDNA elements with shared genomic regions. This may improve the resolution to study ecDNA structural intra/inter-tumor heterogeneity from bulk sequencing.

## 2 Results

### 2.1 An overview of the Decoil algorithm

Decoil reconstructs complex ecDNA structures from long-read nanopore sequencing data using aligned sequencing reads, structural variants (SVs) and coverage profiles as input (Figure 1a). The genome is initially fragmented using a clean breakpoints set (Figure 1a #1). A weighted undirected multigraph is build to encode the structural rearrangements, where nodes are defined as genomic non-overlapping segments and edges represented the structural variants (Figure 1a #2).

**Figure 1.**
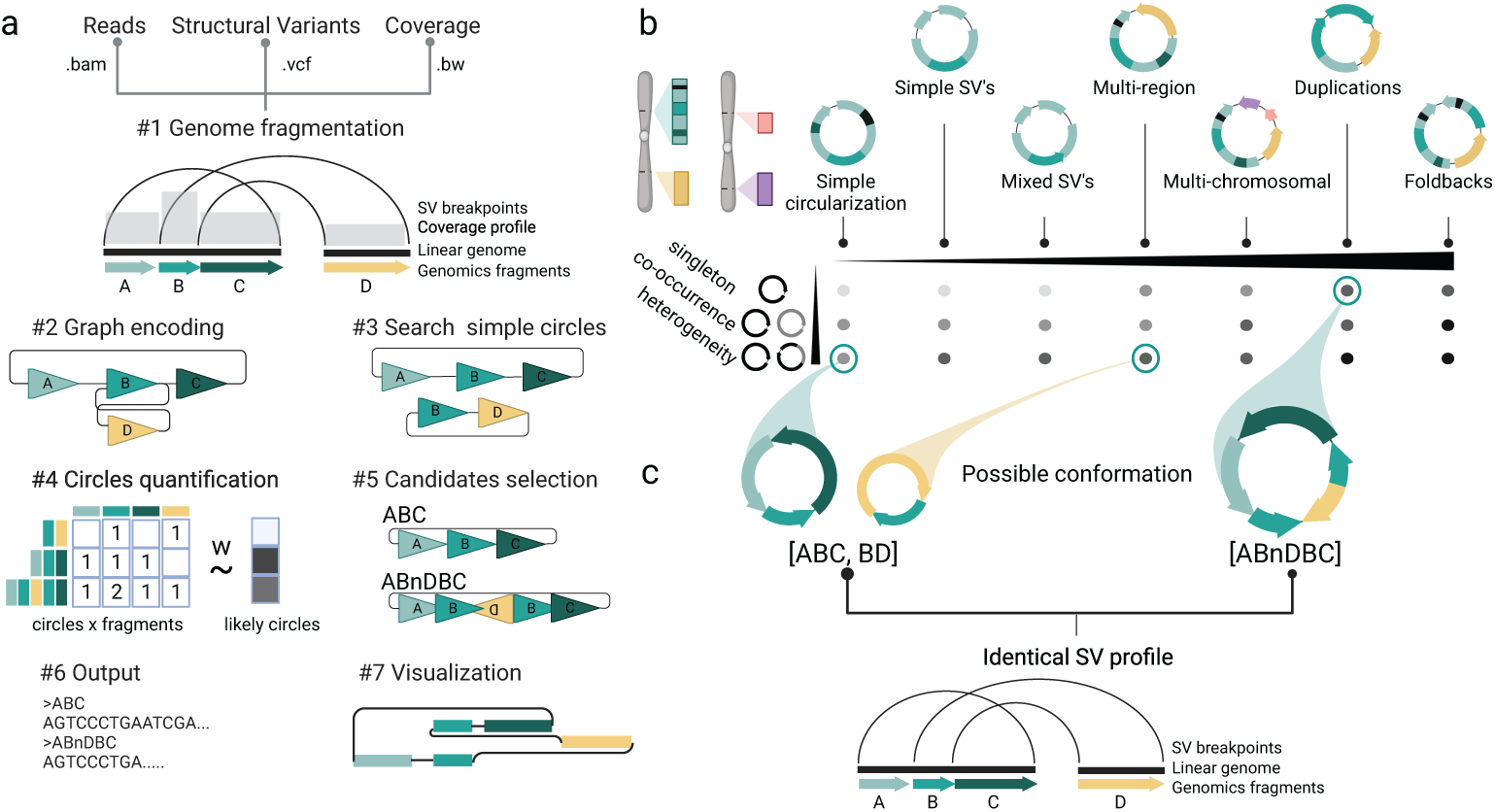
Decoil algorithm overview. (a) Schematic of the Decoil algorithm depicting the major steps (#1 - Genome fragmentation, #2 - Graph encoding, #3 - Search simple circles, #4 - Circles quantification, #5 - Candidates selection, #6 - Output and #7 - Visualization). (b) The overlapping cycles challenge. The left panel displays a heterogeneity scenario, where two different ecDNA elements share a genomic footprint (B fragment), the right panel displays a large structure containing interspersed duplication rearrangement. nD - annotates inverted D fragment. Both scenarios lead to the same SV breakpoint profile.

Next, the graph is explored using a depth-first search approach to discover genome-wide simple circular paths (Figure 1a #3). These can represent a unique circular element or be a sub-component of a more complex circular structure. Subsequently, to account for circular elements containing nested circles, simple circular paths with at least one overlapping genomic fragment are merged into a derived larger circular structure. To avoid exponential growth for the cycles merge, only cycles sufficiently dissimilar are candidates are considered. In order to identify the likely ecDNA elements present in the sample, all simple and derived circle candidates are leveraged as features to fit a *LASSO* regression against the read-alignment mean coverage profile.

This model will (1) select the likely circles explaining the amplification and (2) estimate their proportions within the sample (Figure 1a #4). Using this approach, Decoil can account for ecDNA structures with overlapping genomic footprints (Figure 1b). Lastly, a filtered confident set of circular paths is generated (Figure 1a #5), together with the annotated topology (as defined below), proportion estimates and reconstruction thread visualization (Figure 1a (#6+#7)).

### 2.2 Ranking and simulating ecDNA topologies to capture ecDNA structure diversity

Currently, no guidelines exist for the assessment of ecDNA reconstruction performance from long-read data, nor do benchmarks exist like those for single nucleotide variant (SNV), insertion-deletion (INDEL) and structural variant (SV) detection [16, 17]. The lack of a gold standard datasets for assessing ecDNA reconstruction makes the evaluation of Decoil contingent on high-quality simulated data. The SV profile obtained from the read-alignment was used as information to systematically rank ecDNA by the structure computational complexity. Thus, based on the different SV’s combinations present on the ecDNA element, we propose seven ecDNA topologies (Figure 2): i. Simple circularization, ii. Simple SV’s, iii. Mixed SV’s, iv. Multi-region, v. Multichromosomal, vi. Duplications and vii. Foldbacks. These ecDNA topologies were leveraged to simulate rearrangements on the amplicon in order to create a representative and comprehensive collection of more than 2000 ecDNA templates (Figure 2a), based on which we generated *in-silico* long-read reads at different depths of coverage. This collection serves as a benchmark dataset for evaluating Decoil’s reconstruction performance across varying computational complexities and could be a useful dataset for future ecDNA genomic studies.

**Figure 2.**
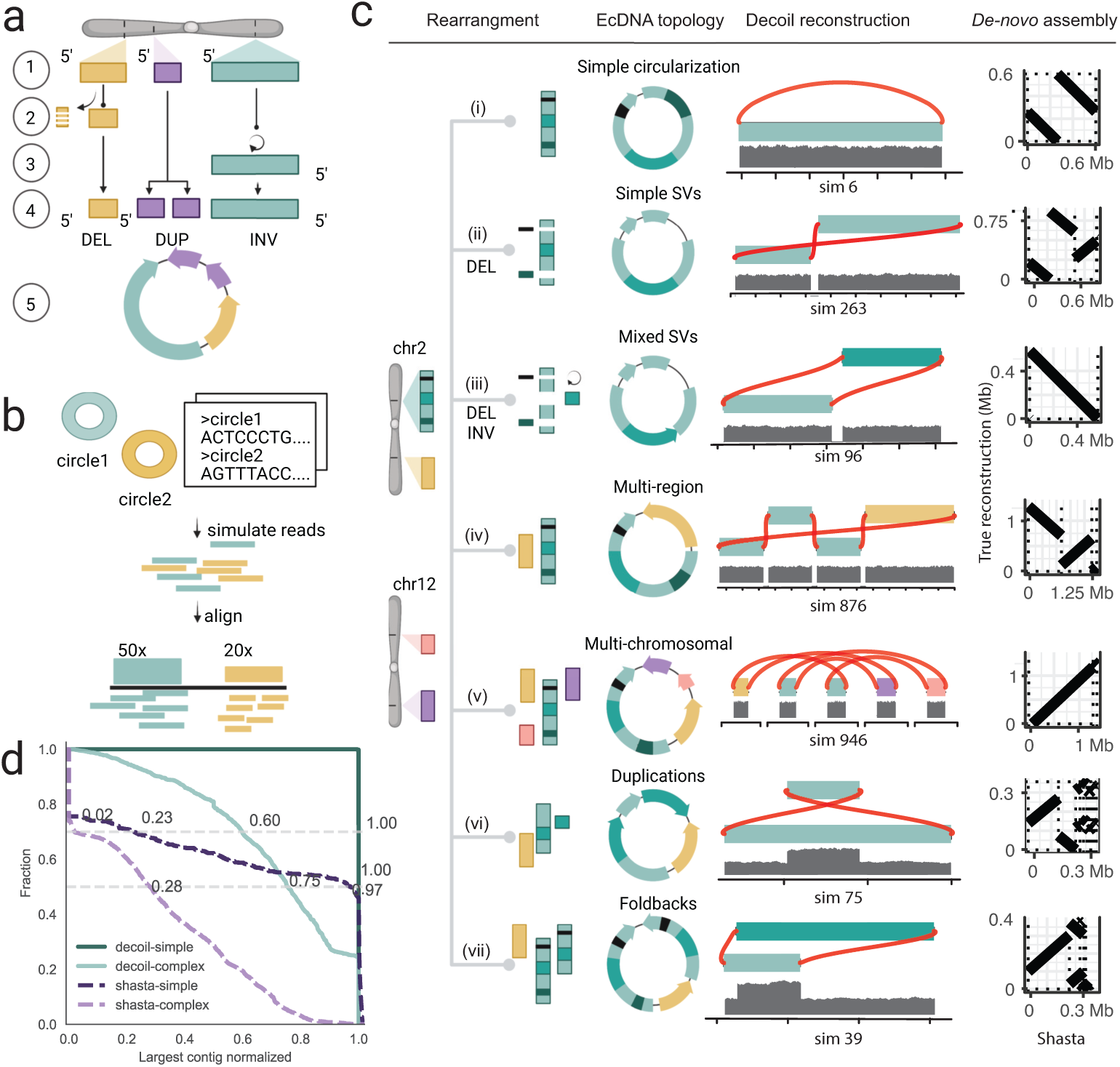
Decoil reconstructs complex ecDNA elements with high fidelity from simulated data. (a) Simulation strategy for individual ecDNA templates generation, describing the main steps: #1 - choose genomic position, #2 - simulate small deletions (DELs), #3 - simulate inversion (INV), #4 - simulate tandem-duplication (DUP), #5 - generate DNA sequence template. The example depicts an ecDNA template harboring 1xDEL (yellow), 1xDUP (purple), and 1xINV (green). (b) *In-silico* long-reads simulation pipeline, based on one or more ecDNA templates, at different depths of coverage. (c) The ecDNA topologies. Reconstruction example by Decoil for a simulated ecDNA element + coverage track. The right column shows the Shastade-*novo* assembly (Y-axis) against the true structure (Y-axis). (d) Decoil and Shasta assembly contiguity for simple (i, ii, iii, iv and v) and complex topologies (vi and vii). X-axis represents the larger contig normalized by the true structure length (1 - a good reconstruction, 0 - poor reconstruction, values > 1 refer to reconstructions larger than the true structure) and Y-axis shows the fraction of reconstructions with the specific contiguity. The gray horizontal lines are at 0.5 and 0.7 fraction.

### 2.3 Decoil’s performance evaluation to reconstruct ecDNA elements in simulated data

The accuracy of ecDNA reconstructions was quantified using the normalized largest contig as a score to measure the assembly contiguity (Section 4.4). Decoil reconstructed simple ecDNA topologies with high-fidelity from simulated data, i.e topologies i, ii, iii, iv and v (> 700 simulations, Figure 2c,d). For the complex topologies, i.e. vi and vii, Decoil reconstructed correctly at least 60% of the true structure (largest contig normalized > 0.6, Figure 2d) in more than 70% of the simulations (> 1900 simulations). Poorly resolved ecDNA elements (largest contig normalized < 0.6) often contained mixed rearrangements including nested duplications and foldbacks, suggesting that such ecDNA elements are more challenging to reconstruct. To demonstrate the utility and feasibility of the method, Decoil was compared against Shasta [18] *de-novo* assembler and CReSIL [15] (Suppl. Figure S7, Suppl. Table S2) using different Quast metrics, (e.g. largest contig, largest alignment, auN) defined in the Extended Methods. CReSIL reconstructs a continuous full alignment for more than 65% of simple topologies with high fidelity (Suppl. Figure S7e). Decoil outperformed Shasta and CReSIL for both, simple and complex topologies in terms of sequence contiguity and completeness (Suppl. Table S1).

### 2.4 Decoil recapitulates ecDNA complexity and their co-occurrence in well characterized cancer cell lines

To show the versatility of the algorithm, Decoil was applied to shallow whole-genome nanopore sequencing of three neuroblastoma cell lines, i.e. CHP212, STA-NB-10DM and TR14, for which ecDNA elements were previously characterized based on various circular DNA enrichment methods and/or validated using fluorescence in situ hybridization (FISH)[6, 19, 20]. Decoil’s reconstructions recapitulated the previously validated ecDNA element in CHP212 with high fidelity (Suppl. Figure S1a,b). An ecDNA harboring *MYCN* and a gene fusion between *SMC6* and *FAM49A* was previously observed in STA-NB-10DM cells [19], which was confirmed by Decoil’s reconstruction (Figure 3a). The ecDNA element in STA-NB-10DM was predicted to be 2.1 MB in size, with an estimated proportions of 171 amplicon copies, harboring an interspersed duplication according to Decoil reconstruction (Figure 3a). Multiple co-occurring ecDNA elements, referred to as ecDNA species in a previous report, were observed in TR14 cells [20]. The three different ecDNA elements, containing *MYCN*, *ODC1* and *MDM2* were reconstructed by Decoil with high fidelity in TR14 (Figure 3b). Additionally, Decoil identified a previously unreported 1.09 MB (Suppl. Table S3), multi-chromosomal ecDNA element containing fragments from chromosome 1 and 2, with an estimated proportion of 22 amplicon copies, harboring *SMC6* and *GEN1* (Figure 3b). This is the largest amplicon and has the lowest number of estimated copies relative to the other co-occurring ecDNA elements, which may be the reason why other reports have not been able to identify it so far. For comparison, the reconstruction’s contiguity in cell lines was evaluated also using Shasta. For CHP212, the agreement between Decoil and Shasta [18] was 100% (Suppl. Figure S1b,c). In STA-NB-10DM, the interspersed duplication on the ecDNA indicates increasing reconstruction complexity. Thus, Shasta did not assemble a contiguous circular element (Suppl. Figure S2a), whereas Decoil identified a contiguous circular path through the graph of this ecDNA element (Figure 3a). For TR14, the structures of amplicons harboring *SMC6*, *MDM2* or *ODC1* were consistent between Decoil and Shasta (Suppl. Figure S3, Suppl. Figure S2b). Additionally, the *MYCN* - containing ecDNA was reconstructed by Decoil (Figure 2b), but was not fully resolved by Shasta (Suppl. Figure S4) due to overlapping rearrangements at the *MYCN* locus (Suppl. Figure S2b). Thus, Decoil is a versatile algorithm to (1) reconstruct complex ecDNA elements in cancer cell lines and (2) discover previously unknown ecDNAs from long-read sequencing data.

**Figure 3.**
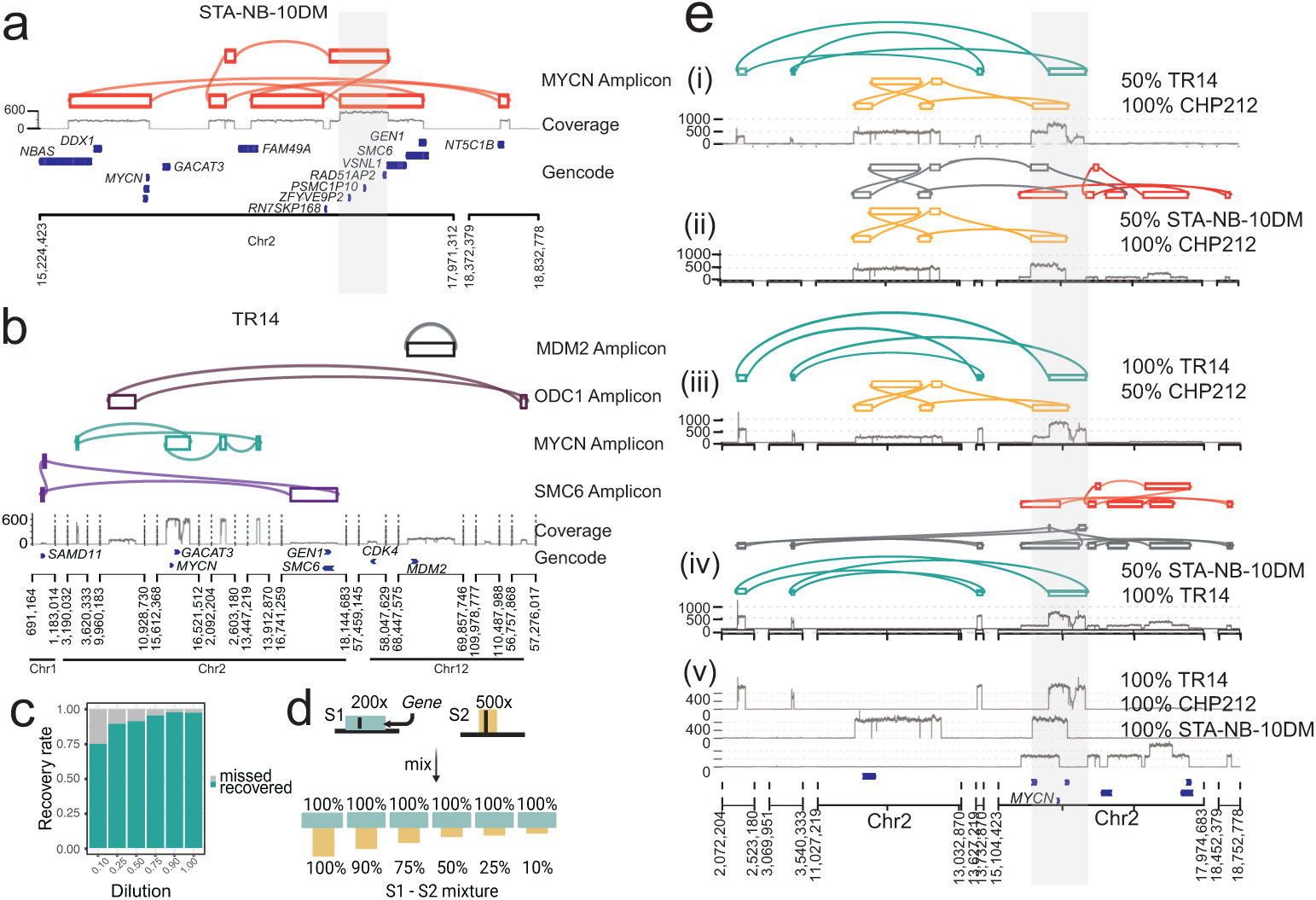
Decoil captures the ecDNA structure complexity and heterogeneity in neuroblastoma cell lines. (a) STA-NB-10DM amplicon reconstruction by Decoil (top), coverage track (middle) of the aligned reads to reference genome GRCh38/hg38 and GENCODE v42 annotation (bottom). The grey highlighted region *chr*2 : 17221081 17538185 (GRCh38/hg38) is an interspersed duplication. (b) TR14 amplicons co-occurrence reconstructed by Decoil (top four tracks), the coverage track (middle) and GENCODE V42 annotation (bottom). (c) ecDNA breakpoints recall (Y-axis) for *in-silico* ecDNA mixtures, by dilutions (Y-axis). The *MYCN*-amplicon for CHP212, TR14 and STA-NB-10DM is composed of 10, 8 and 14 breakpoints, respectively. For TR14 all ecDNA breakpoints are considered from amplicons originating from *MYCN*, *ODC1* (4 breakpoints), *MDM2* (2 breakpoints) and *SMC6* (6 breakpoints). (d) *In-silico* dilution strategy, where two samples S1 (green) and S2 (yellow) are mixed at different ratios to generate ecDNA mixtures. (e) EcDNA reconstruction by Decoil for *in-silico* ecDNA mixtures of cell lines. (i to iv) display reconstructed overlapping ecDNA elements harboring *MYCN* (grey highlight) in different mixtures, originating from TR14 (green), CHP212 (yellow) or STA-NB-10DM (orange). (v) Coverage track for pure TR14, CHP212 and STA-NB-10DM samples. In grey misassemblies are depicted.

**Figure 4.**
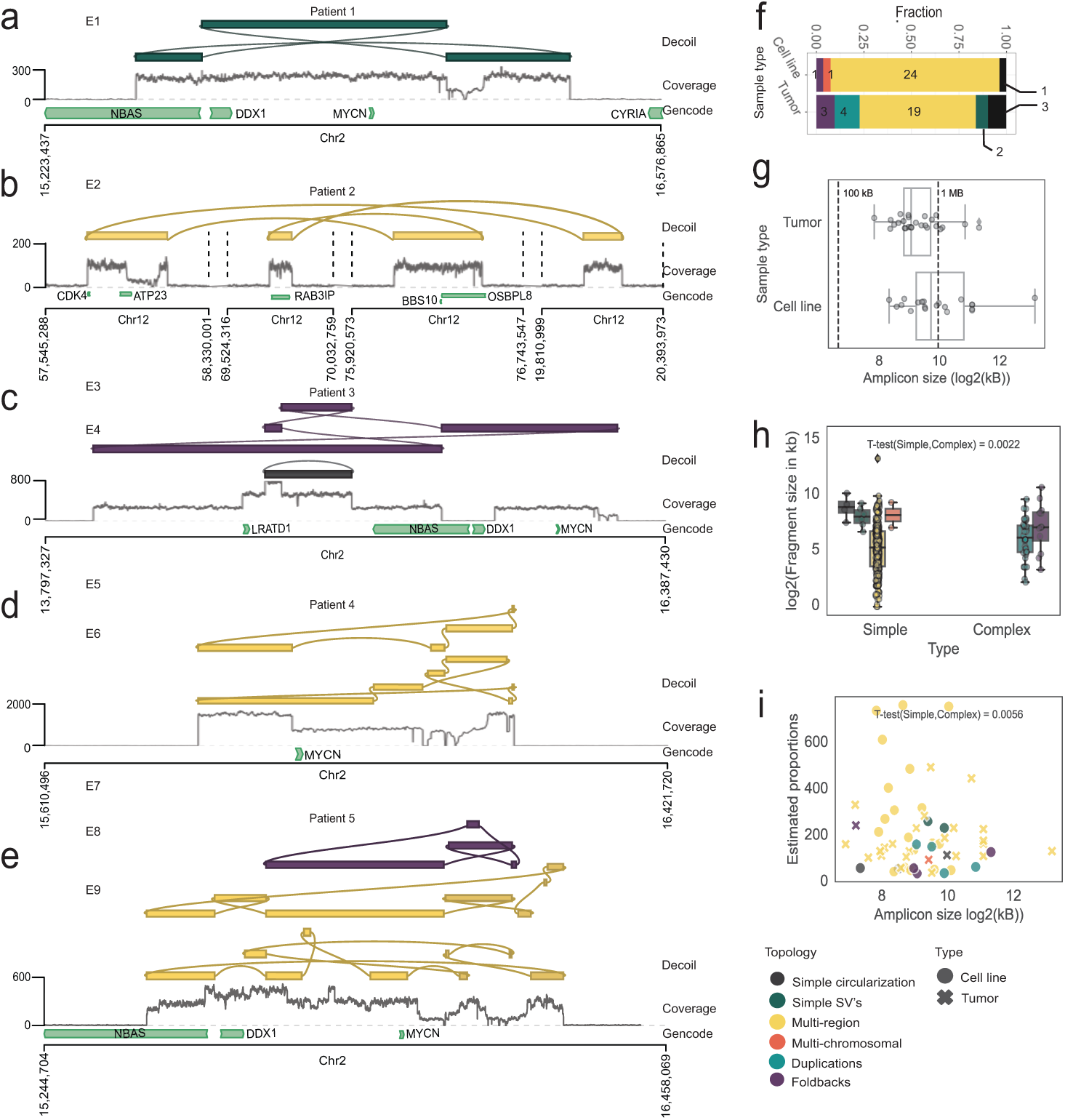
Decoil recovers structurally complex ecDNA elements in primary cancers. Examples of ecDNA structure reconstruction of (a) Simple SV’s, (b,d) Multi-region, (c) Foldbacks and (e) Duplication/Foldbacks topologies in patient samples. For (a-e) from top to bottom are the Decoil reconstruction, nanopore coverage of the aligned reads to reference genome GRCh38/hg38 and GENCODE v42 annotation tracks. Top 3 reconstruction per sample, labeled as ecDNA, with an estimated proportions >= 30 copies were included. E1-E9 are the ids for each reconstruction (Suppl. Table S4). (f) Decoil ecDNA reconstructions topology spectrum in 5 cell lines and 9 patient samples. (g) Amplicon size (Y-axis) distribution (Y-axis) for all data (5 cell lines, 9 primary tumor samples). (h) ecDNA fragment size distribution split for simple (Simple circularization, Simple SV’s, Multi-region, Multi-chromosomal) or complex (Duplications, Foldbacks) topologies. T-test statistics was applied for the the fragment size of simple and complex topologies. (i) Amplicon size (Y-axis) against estimated proportions (Y-axis) displayed by Decoil. T-test statistics was applied for the estimated proportions of simple and complex topologies. Amplicons labeled as ecDNA and with an estimated proportions >= 30 copies were included in (f,g,h,i). Boxplot shows Q1(25%), Q2(median) and Q3(75%), interquartile range IQR = Q3 - Q1, and whiskers are 1.5 x IQR. Colors in (a-f, h) correspond to the legend in panel (i).

### 2.5 Decoil can recover ecDNA structure heterogeneity

To demonstrate that Decoil can resolve structurally distinct ecDNA elements with overlapping genomic footprint, we generated 33 *in-silico* mixtures, by pair-wise combination of three neuroblastoma cell lines at different ratios, i.e. CHP212, STANB-10DM and TR14, each containing a structurally distinct ecDNA element haboring *MYCN* gene (Figure 3d, Section 4.5). The breakpoint junctions of the individual ecDNA elements were recovered in the different mixtures with a recall of 93% (Figure 3c). These results suggest that Decoil can distinguish between different co-occurring ecDNA elements with overlapping genomic footprints, enabling the measurement of structural ecDNA heterogeneity.

### 2.6 Exploring structural ecDNA complexity in cancer patients using Decoil

In order to explore structural ecDNA complexity in tumors, shallow whole-genome nanopore sequencing on a cohort of 13 neuroblastomas was performed, of which ten harbored at least one ecDNA (experimentally confirmed by FISH) and three negative controls (no ecDNA present). Decoil did not detect any ecDNA in the negative control cohort and reconstructed at least one amplicon for the other 9 samples, with genomic fragments originated from chromosome 2 or chromosome 12. The reconstructed ecDNA elements varied greatly in their complexity (Figure 4f) and ranged from very simple (Figure 4a) or multi-region (Figure 4b) to heavily rearranged complex structures (Figure 4c,d,e). Decoil reconstructs for Patient 4 two ecDNA elements with an individual estimated proportions of more than 700x, resolving the same breakpoints as previously published ([8], Fig. 7a). For same patients Decoil reconstructed multiple circular elements with different estimated relative proportions, which suggests ecDNA structural heterogeneity (Figure 4e). Multi-region topology seemed to be the most frequent ecDNA topology identified in patients, consistent with the ecDNA elements detected in cell lines (Figure 4f). Decoil reconstructed ecDNA elements with a mean size of 1.4 MB in cell lines and 0.7 MB in patient samples (Figure 4g), in line with other studies [5]. Contiguous genomic fragments on ecDNA had a mean size of 127 kb in cell lines and 145 kb in patient samples (Suppl. Figure S5b). While the ecDNA size was conserved for the different topologies (Suppl. Figure S5a), complex ecDNA elements had significantly shorter fragments than simple ecDNAs (Figure 4h, Suppl. Figure S5c). Lastly, simple ecDNA had higher copy numbers than complex ones in this cohort (Figure 4i, Suppl. Figure S5d) and may indicate yet unknown structural features that may influence ecDNA maintenance and/or oncogene regulation.

### 2.7 Scalability and runtime

Using the simulated (0.01 WGS mean coverage) and real dataset (3-7X WGS mean coverage) we show a runtime of Decoil in the range of minutes or less per sample for Decoil standalone and several hours for Decoil-pipeline, with a maximum memory usage of less than 5 GB, on 4 x threads (Suppl. Figure S8).

## 3 Discussion

The structural complexity and heterogeneity of ecDNA make its reconstruction from sequencing data a challenging computational problem. We here presented Decoil, a method to reconstruct co-occurring complex ecDNA elements.

Due to their random mitotic segregation, many ecDNA elements, which may structurally differ, co-occur in the same cancer cells [8]. Disentangling ecDNA with shared genomic regions has not yet been addressed by other methods, and it cannot be resolved by *de-novo* assemblers (e.g. Shasta) when sequencing reads are smaller than the size of genomic fragments (mean length > 125 kb in our cohort) within an ecDNA element. Decoil uses *LASSO* regression to reconstruct distinct ecDNA elements with overlapping genomic footprint, which enables the exploration of ecDNA structural heterogeneity. We have chosen this approach as it performed reasonably in our hands compared to other linear regression models (Suppl. Figure S6). One limitation of our methods represent the correct decomposition into distinct ecDNA elements for structures containing repetitive regions. This would lead to incomplete structural resolution, e.g. the order of the repeat-containing genomic segments might remain ambiguous. Furthermore, ecDNA present at low abundance or SVs not detected due to computational limits may affect Decoil’s performance. Measuring the limit of detection of Decoil was not addressed in this manuscript, as it will require comprehensive tumor datasets with validated ecDNA structures. Ultra-long read sequencing (>100 kb) at high coverage, or other sequencing technologies, may improve the SV detection and structural resolution of ecDNA using Decoil, but aforementioned scenarios may remain difficult to resolve.

A structure-function relationship was first demonstrated for ecDNA by reports describing regulatory elements on ecDNA [6, 10, 20, 21]. These reports revealed that complex ecDNA rewire tissue-specific enhancer elements to sustain high oncogene expression [6, 22]. This also occurs through formation of new topologically associated domains [6]. Decoil was able to identify multi-region ecDNA elements, which were previously linked to enhancer hijacking [6], suggesting that it may help map such alterations in cancer. We envision that combining Decoil with DNA methylation analysis from the same nanopore sequencing reads may enable exploration of potential regulatory heterogeneity in co-occurring ecDNA elements, which was not previously possible.

The reconstruction of ecDNA in a cohort of neuroblastoma tumors and cell lines using Decoil suggested that structurally simple ecDNA elements occurred at higher copy numbers and were larger in size compared to complex ecDNA. This might be due to computational biases, as complex structures are more difficult to reconstruct, and certainly needs to be verified in larger tumor cohorts. However, it is reasonable to speculate that ecDNA complexity could influence ecDNA maintenance or impact its copy number in yet unidentified ways. Future analyzes using Decoil may help verify this observation and address such questions.

In summary, we envision that Decoil will advance the exploration of ecDNA structural heterogeneity in cancer and beyond, which is essential to better understand mechanisms of ecDNA formation and its structural evolution and may serve as the basis to identify DNA elements required for oncogene regulation and ecDNA maintenance.

## 4 Methods

### 4.1 Decoil algorithm

Decoil (<u>d</u>econvolve <u>e</u>xtrachromosomal <u>c</u>ircular DNA isoforms from long-read data) is a graph-based method to reconstruct circular DNA variants from shallow long-read WGS data. This uses (1) structural variants (SV) and (2) focal amplification information to reconstruct circular ecDNA elements. The algorithm consists of six modules: genome fragmentation, graph encoding, search simple circles, circles quantification, candidates selection, output, and visualization.

#### Genome fragmentation

The SVs are filtered based on multiple criteria. Only SVs flagged as ’PASS’ or ’STRANDBIAS’, having on target coverage >= 5X (default) and VAF (Variant Allele Frequency) >= 0.2 (default) are kept. Breakpoints in a window size of 50 bp are merged. This curated breakpoints set *s* is used to segment the genome into *n* + 1 non-overlapping fragments *f F*, where *F* represents the non-overlapping fragments set.

#### Graph encoding

The coverage profile, read alignment data and fragments set *F* are combined to build a weighted undirected multigraph, denoted as *G* = (*V, E*). In *G*, a vertex *f* represents a genomic fragment from the set *F*, and an edge *e* represents a SV connecting two fragments. A multigraph is used to represent scenarios where multiple SVs share the same breakpoints. The vertices *f* in the graph *G* are objects, each consisting of two internal nodes: (1) a ’head’ node and (2) a ’tail’ node, which are used to track the orientation of genomic fragments. The edges have two properties, (1) length defined as the SV length and (2) weight defined as *DR* (coverage of alternative variant). The SVs are encoded in the graph *G* based on their annotated type:

- BND, DEL - one edge connects head to tail of the two fragments
- DUP - one edge connects tail to head of the two fragments
- INV, INVDUP - two edges connect head to head and tail to tail of the two fragments
- Fragments with a mean coverage <= 5X (default) or standalone (*degree*(*v*) = 0) are discarded from the graph.

#### Search simple circles

Decoil continues by searching all simple circular paths *c* in the graph *G* using weighted depth-first search (DFS) approach. A cycle in a DFS tree is defined as a path where two visited nodes, *u* and *v*, are connected through a backedge (*u, v*), with *u* being the ancestor of *v*. This approach conducts a genome-wide search for circular paths and it traverses the tree in descending sorted order of the edges to ensure consistent results. The identified cycles are hashed and saved in a canonical form, where the leftmost fragment corresponds to the smaller genomic position. Duplicated cycles are removed during tree exploration. The resulting set comprises *S* unique simple cycles, allowing for shared sub-paths. These simple cycles, denoted as *c S*, are then organized into non-overlapping clusters (*M*), where two cycles *S* belong to same cluster if they share at least one genomic fragment.

#### Circles quantification

A set of derived cycles (*D*) was created by performing all the combinations between all simple cycles *c M*. The reasoning behind it is to allow reconstruction of complex structures, e.g. containing large duplications. To distinguish between true possible circular ecDNA elements and artifacts a *LASSO* regression is fitted against targets *Y* ^|^*^F^* ^|^ using input *X*^|^*^F^* ^|⇥|^*^S^*^||^*^D^*^|^, to learn the proportions of the cycles. *x_jik_ 2 X* is defined as the occurrence of fragment *f_jk_* in circle *c_ik_* and *y_jk_ 2 Y* represents the total mean coverage spanning fragment *j*, belonging to cluster *m_k_*. *LASSO* was performed for each cluster *m_k_*.

*LASSO* regression definition

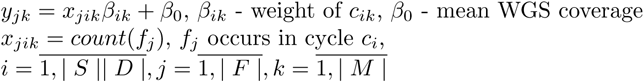

After *LASSO* regression fit, circles *c_ik_* with weight *β_ik_ > t* were kept, where threshold *t* = *max*(*min*(*coverage*(*f_j_*))*/*4, 10). The obtained *LASSO* coefficient *β_ik_* represent the estimated proportions of each cycle *c_ik_*. The higher the value the more likely is the cycle to be a true ecDNA element. To avoid overfitting of the model, a penalty term *alpha* = 0.1 was used.

#### Candidates selection

From the candidates list we filter out cycles with an estimated proportions <= mean WGS coverage (default). Lastly, circular elements larger than 0.1 MB (threshold published by [13]) are labeled as ecDNA.

#### Output and Visualization

The algorithm outputs the candidates list as *.bed, *.fasta, including the mean coverage and orientation per fragment, estimated proportions of circular element and the annotated topology (as defined in the paper). The *summary.txt* displays all found circular elements. Lastly, visualization of the individual ecDNA reconstructions threads are generated based on gGnome R package (https://github.com/mskilab/gGnome). A *.html report is generated to summarize all ecDNA reconstruction threads found in the sample.

### 4.2 Ranking ecDNA topologies definition

To assess Decoil’s reconstruction performance, we generated an *in-silico* collection of ecDNA elements, spanning various sequence complexities for systematic evaluation. We introduced a ranking system and defined seven topologies of increasing computational complexity, based on the SV’s contained on the ecDNA element: (1) *Simple circularization* - no structural variants on the ecDNA template, (2) *Simple SV’s* - ecDNA contains either a series of inversions or deletions, (3) *Mixed SV’s* - ecDNA has a combination of inversions and deletions, (4) *Multi-region* - ecDNA contains different genomic regions from the same chromosome (DEL, INV and TRA allowed), (5) *Multi-chromosomal* - ecDNA originates from multiple chromosomes (DEL, INV and TRA allowed), (6) *Duplications* - ecDNA contains duplications defined as a region larger than 50 bp repeated on the amplicon (DUP’s + other simple rearrangements), (7) *Foldbacks* - ecDNA contains a foldback defined as a two consecutive fragments which overlap in the genomic space, with different orientations (INVDUP’s + all other simple SV’s). Every topology can contain a mixture of all other low-rank topologies.

### 4.3 Simulate ecDNA

The simulation framework contains probabilistic variables, which model the chromosome weights, fragment position, fragment length, small deletion ratio, inversion ratio, foldback ratio, and tandem-duplication ratio. To cover a wide range of possible conformations more than 2000 ecDNA sequence templates were generated. Based on these definitions, *in-silico* ecDNA-containing samples were generated by simulating noisy long-reads, at different depth of coverage, with an adapted version of PBSIM2 (Ono et al. 2021 [23]). This workflow is available under https://github.com/madagiurgiu25/ecDNA-simulate-validate-pipeline. See Extended Methods for detailed description.

### 4.4 Performance evaluation on simulated data

To evaluate the correctness of reconstruction for Decoil, Shasta and CReSIL, Quast [24] 5.2.0 was applied to compute different metrics (https://quast.sourceforge.net/docs/manual.html). To overall reconstruction performance was quantified as the mean and standard deviation of the largest contig metric. See Extended Methods for detailed description.

### 4.5 Evaluate amplicon’s breakpoints recovery in ecDNA mixtures

To evaluate how well we reconstruct amplicons with overlapping footprints we generate a series of dilutions by mixing the CHP212, STA-NB-10DM and TR14 cell lines at different ratios. We generated two types of mixtures. First, we combine 100% of one sample with different percentages of another sample, i.e. 10, 25, 50, 75, 90, 100% (Figure 3c). Secondly, we generate mixtures at different ratios for both samples (10-90, 25-75, 50-50, 75-25, 90-10%). Picard 2.26 (https://broadinstitute.github.io/picard/) was used to downsample the .bam file to 10, 25, 50, 75, 90% and samtools 1.9 to merge the different ratios to create *in-silico* ecDNA mixture. SV calling was performed using sniffles [25] 1.0.12, followed by Decoil reconstruciton. The reconstructed ecDNA elements in mixtures were evaluated using as metric the breakpoints recall, defined as the ratio of true breakpoints found in mixtures.

### 4.6 Runtime and memory benchmarking

For both simulated and real datasets, we conducted an analysis of the runtime and memory usage. The runtime, including the raw elapsed time (ElapsedRaw) and CPU time (CPUTime), was measured. Additionally, memory usage was assessed using the maximum resident set size (MaxRss). These metrics were derived from the Slurm output, providing insights into the computational resources consumed during the analysis.

## 5 Data access

The cell lines data and patient data is available under EGAXXXXX and accessible upon reasonable request.

## 6 Code availability

Decoil is available freely as docker container. https://github.com/madagiurgiu25/decoilpre. It can be run in two different ways: (1) decoil-pipeline, a user-friendly snakemake-workflow [26] which takes as input a .bam file and computes internally the SV calling, the coverage profile, ecDNA reconstruction and visualization, (2) decoil standalone, for more advanced and flexible usage, requires as input a .vcf file with the precomputed SV calling, a .bw file with the coverage profile and a .bam file. With this article we publish other several associated tools for: a ecDNA sequence simulator based on specified topology https://github.com/madagiurgiu25/ecDNA-sim, a long-read ecDNA containing samples simulator (adapted PBSIM2 for circular reference) https://github.com/madagiurgiu25/pbsim2, a snakemake [26] processing and validation pipeline for ecDNA containing simulated samples https://github.com/madagiurgiu25/ecDNA-simulate-validate-pipeline.

## 7 Acknowledgements

We would like to thank Roland F. Schwarz, Julia Markowski, and Svenja Mehringer for their input and thoughtful suggestions during the development of this paper. We thank the Berlin Institute of Health (BIH) team for the support and providing the necessary infrastructure. Computation was performed on the HPC for Research cluster of the BIH. We thank the patients and their parents for granting access to the tumor specimens and clinical information that were analyzed in this study. We thank the Neuroblastoma Biobank and Neuroblastoma Trial Registry (University Children’s Hospital Cologne) of the GPOH for providing samples.

## 8 Declarations

### 8.3 Ethics approval

Patients were registered and treated according to the trial protocols of the German Society of Pediatric Oncology and Hematology (GPOH). This study was conducted in accordance with the World Medical Association Declaration of Helsinki (2013) and good clinical practice; informed consent was obtained from all patients or their guardians. The collection and use of patient specimens was approved by the institutional review boards of Charité-Universitätsmedizin Berlin and the Medical Faculty, University of Cologne. Specimens and clinical data were archived and made available by Charité-Universitätsmedizin Berlin or the National Neuroblastoma Biobank and Neuroblastoma Trial Registry (University Children’s Hospital Cologne) of the GPOH. The *MYCN* gene copy number was determined as a routine diagnostic method using FISH.

## Appendix A Supplementary Figures

**Suppl. Table S1.**
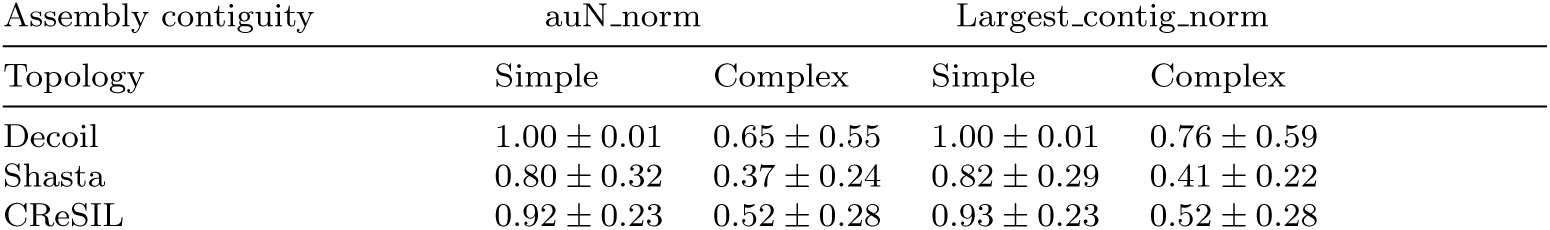
Performance evaluation for Decoil and Shasta. Every value in the table represents the mean and standard deviation for the entire simulated dataset. All the metrics are normalised by the simulations true length. 1 means correct assembly, < 1 assembly regions were missed, > 1 assembly regions were additional included.

**Suppl. Table S2.** Assembly contiguity for Decoil and Shasta in simulated data. The metrics are computed using Quast. All the metrics are normalised by the simulations true length. For Largest contig norm, Total length norm, N50 norm, auN norm, Largest alignment norm (not available for Shasta), Total aligned length norm 1 means correct assembly, *<* 1 assembly regions were missed, *>* 1 assembly regions were additional included. The Misassembled contigs length norm shows how much (percentage) of the assembled output diverge from the reference. Unaligned length norm represents genomic region percentage missed by the reconstruction or assembly (see Supplementary Methods for the full definition of these metrics).

|  | Shasta | Decoil | CReSIL |
| --- | --- | --- | --- |
| Largest_contig_norm | 0.53 ± 0.31 | 0.82 ± 0.54 | 0.74 ± 0.23 |
| Total_length_norm | 0.91 ± 0.99 | 1.03 ± 0.61 | 0.75 ± 0.22 |
| N50_norm | 0.52 ± 0.33 | 0.8 ± 0.54 | 0.74 ± 0.23 |
| N90_norm | 0.43 ± 0.38 | 0.51 ± 0.6 | 0.73 ± 0.24 |
| auN_norm | 0.49 ± 0.33 | 0.74 ± 0.52 | 0.74 ± 0.23 |
| Misassembled_contigs.length_norm | 0.05 ± 0.19 | 0.23 ± 0.37 | 0.36 ± 0.39 |
| Unaligned_length_norm | 0.48 ± 1.09 | 0.02 ± 0.47 | 0.0 ± 0.0 |
| Largest_alignment_norm | / | 0.73 ± 0.29 | 0.56 ± 0.3 |
| Total_aligned_length_norm | 0.49 ± 0.35 | 1.0 ± 0.34 | 0.74 ± 0.22 |

**Suppl. Table S3.** Amplicons overview across cell lines. Size, estimated proportions and topology output by Decoil.

| Cell line | Amplicon | Size MB | Est. prop. | Topology |
| --- | --- | --- | --- | --- |
| CHP212 | *MYCN* | 1.69 | 440 | Multi-region |
| STA-NB-10DM | *MYCN* | 2.1 | 253 | Multi-region |
| TR14 | *SMC6* | 1.09 | 20 | Multi-chromosomal |
| TR14 | *MDM2* | 1.01 | 109 | Simple circularization |
| TR14 | *MYCN* | 0.7 | 487 | Multi-chromosomal |
| TR14 | *ODC1* | 0.68 | 89 | Multi-chromosomal |

**Suppl. Figure S1.**
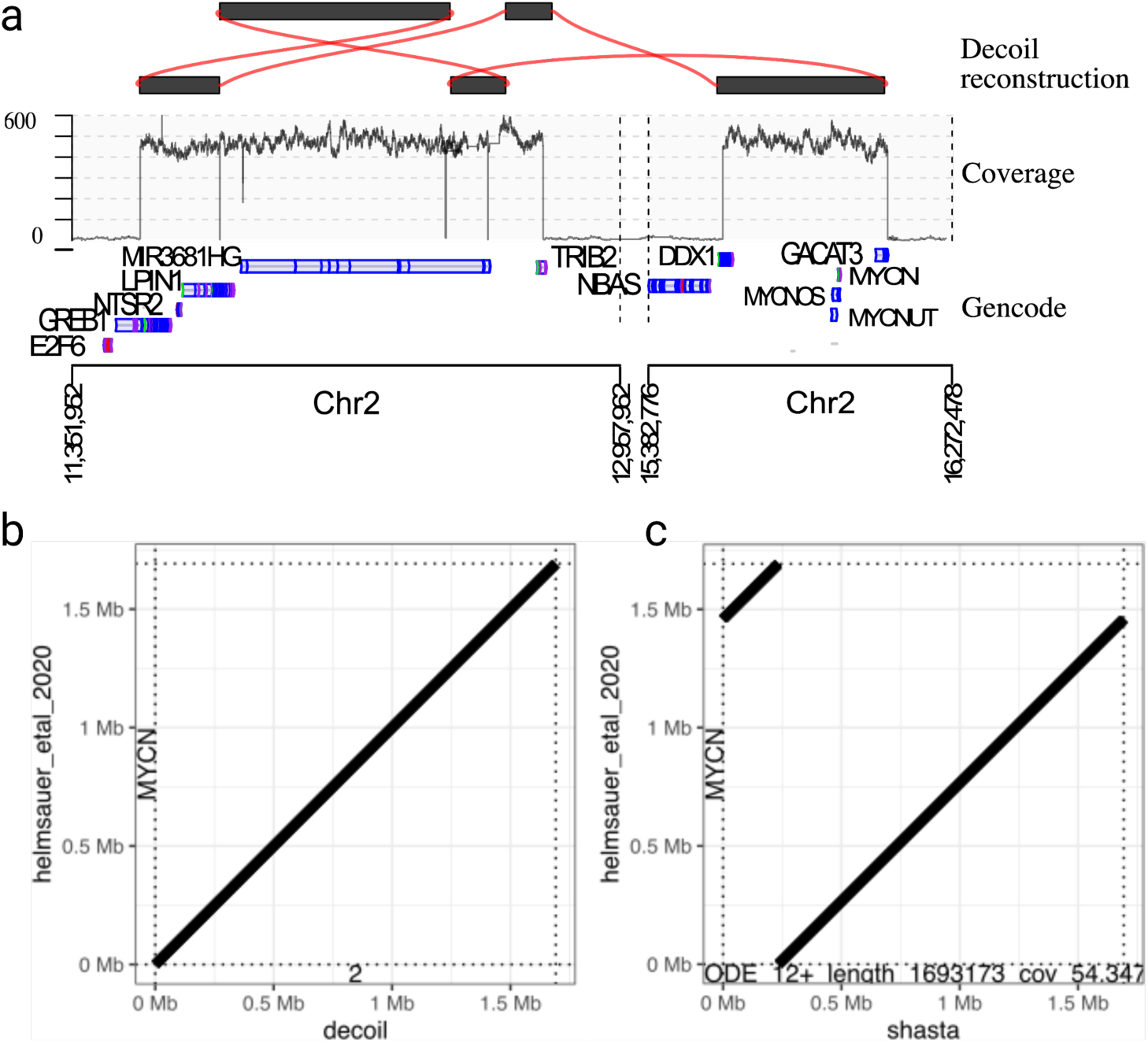
CHP212 amplicon reconstruction. (a) Decoil reconstructs an multi-region intra-chromosomal ecDNA structure (top), for which the read-alignment coverage (middle) and GENCODE v42 gene annotation (bottom) are displayed. (b) Sequence identity comparison between Decoil (X-axis) and published coordinates Helmsauer et al. 2020 (Y-axis). (c) Sequence identity comparison between Shasta (X-axis) and published coordinates Helmsauer et al. 2020 (Y-axis).

**Suppl. Table S4.** EcDNA elements reconstruction description in patients by Decoil.

| Patient | ID | Size MB | Est. prop. | Topology |
| --- | --- | --- | --- | --- |
| Patient1 | E1 | 0.9 | 226 | Simple SV's |
| Patient2 | E2 | 1.09 | 44 | Multi-region |
| Patient3 | E3 | 2.55 | 122 | Foldbacks |
| Patient3 | E4 | 0.3 | 50 | Simple circularization |
| Patient4 | E5 | 0.22 | 732 | Multi-region |
| Patient4 | E6 | 0.39 | 756 | Multi-region |
| Patient5 | E7 | 0.5 | 53 | Foldbacks |
| Patient5 | E8 | 0.77 | 47 | Multi-region |
| Patient5 | E9 | 0.41 | 55 | Multi-region |

**Suppl. Figure S2.**
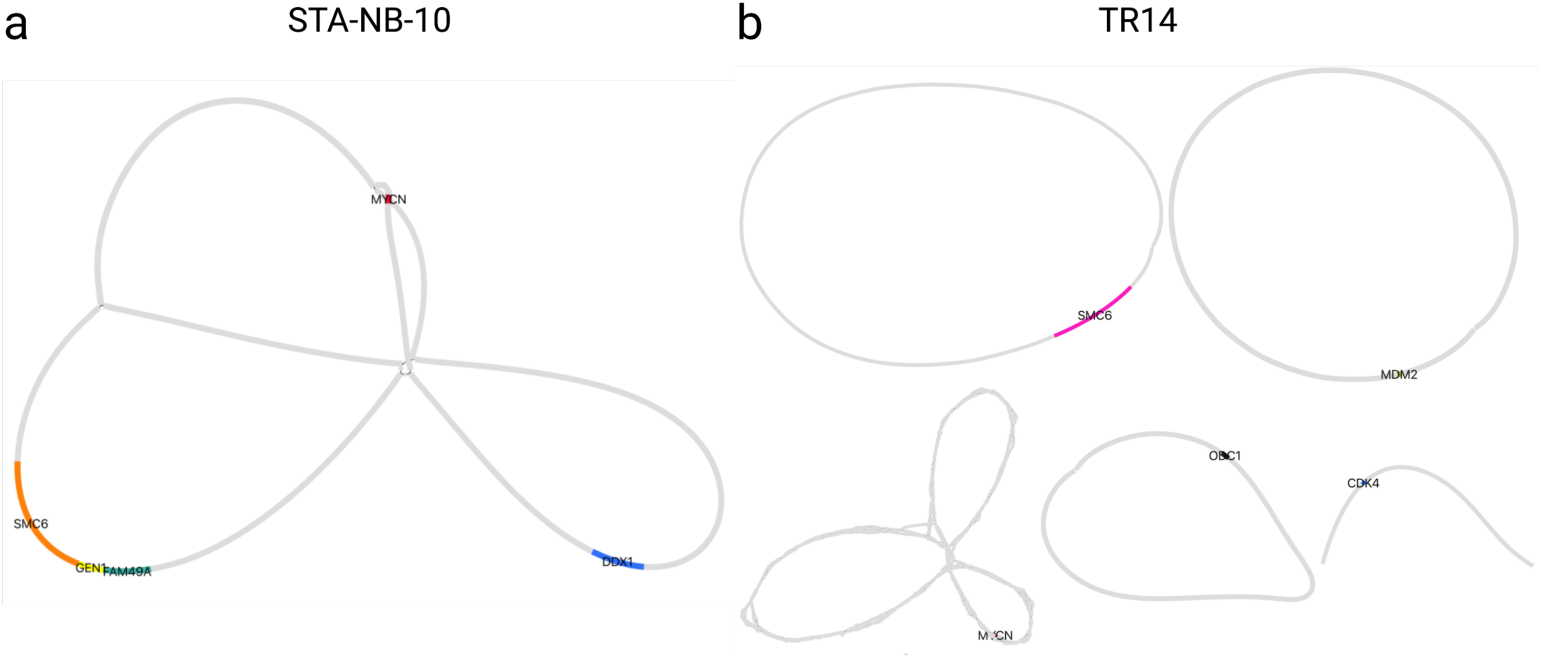
Amplicons de-novo assembly using Shasta. Bandage output showing the assemblies for all found ecDNA elements in two neuroblastoma cell lines. (a) STA-NB-10DM containing one ecDNA element with co-amplification of *MYCN*, *DDX1*, *GEN1* and the fusion *SMC6* - *FAM49A*. Note that *SMC6* overlaps with *GEN1*. (b) TR14 containing four circular assemblies *SMC6*, *ODC1*, *MDM2* and *MYCN*, whereas *CDK4* is resolved as a linear contig. *MYCN* locus is not a contiguous structure.

**Suppl. Figure S3.**
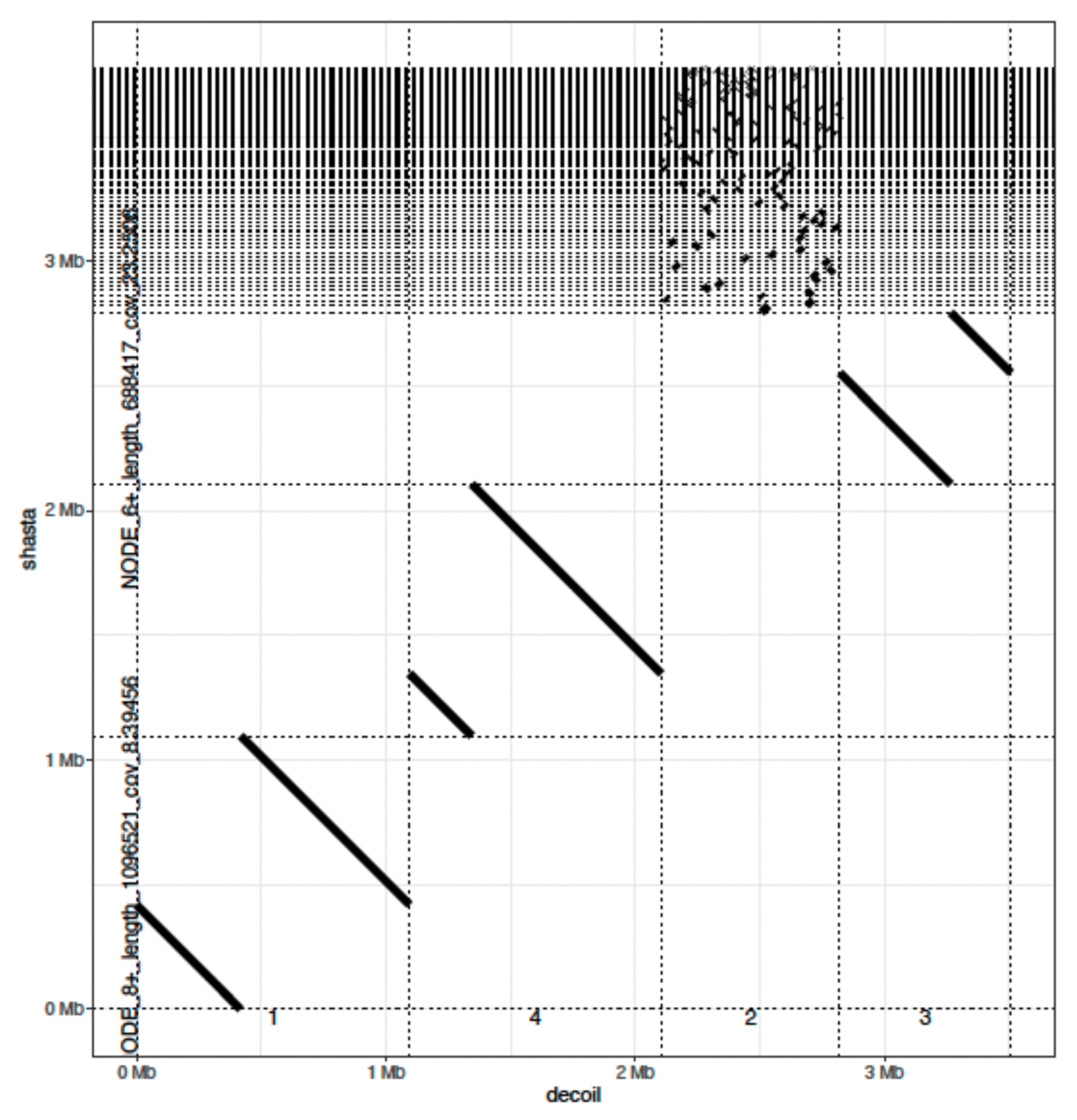
Decoil and Shasta agreement for TR14 ecDNA elements. The dotplot shows the sequence identity between Shasta (Y-axis) and Decoil (X-axis). On the X-axis the numbers mean: 1 - *SMC6*, 4 - *MDM2*, 2 - *MYCN*, 3 - *ODC1* amplicon.

**Suppl. Figure S4.**
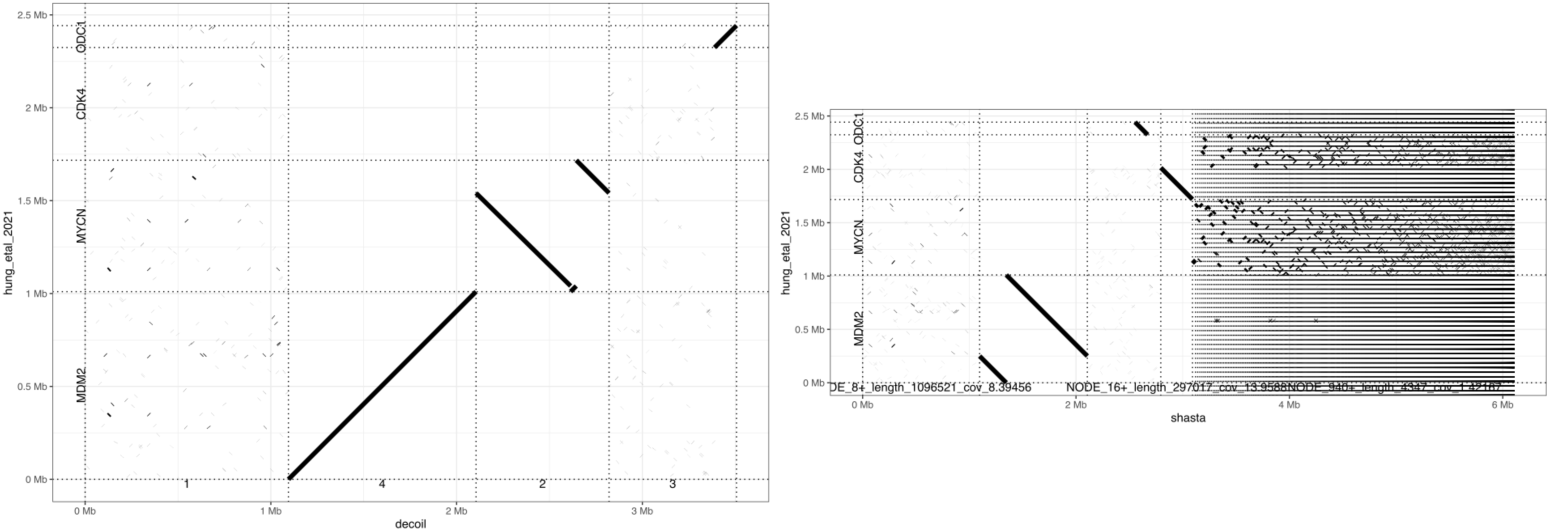
Sequence identity between Decoil and Shasta reconstructions (X-axis) and Hung et al. 2021 (Y-axis) for the TR14 ecDNA elements.

**Suppl. Figure S5.**
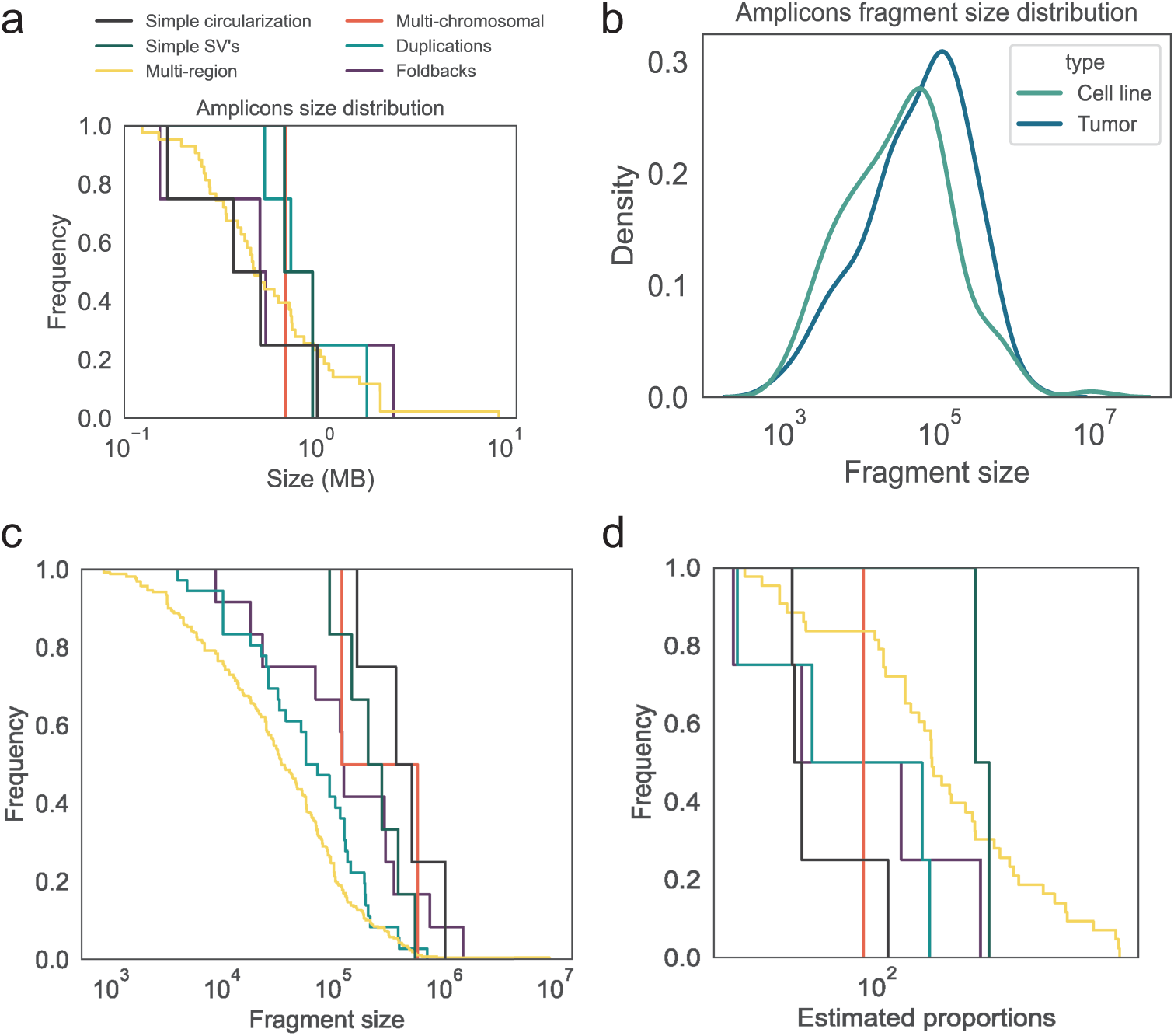
EcDNA amplicon features (extended). (a) Frequency (Y-axis) of the amplicon size (X-axis) for the identified ecDNA topologies across cell lines and patient samples. (b) Fragment size (X-axis) distribution (Y-axis) of the reconstructed amplicons. (c) Fragment size (X-axis) frequency. (d) Frequency of the estimated proportions of the amplicons across the different identified topologies. The colors in (c,d) corresponds to the legend in (a).

**Suppl. Figure S6.**
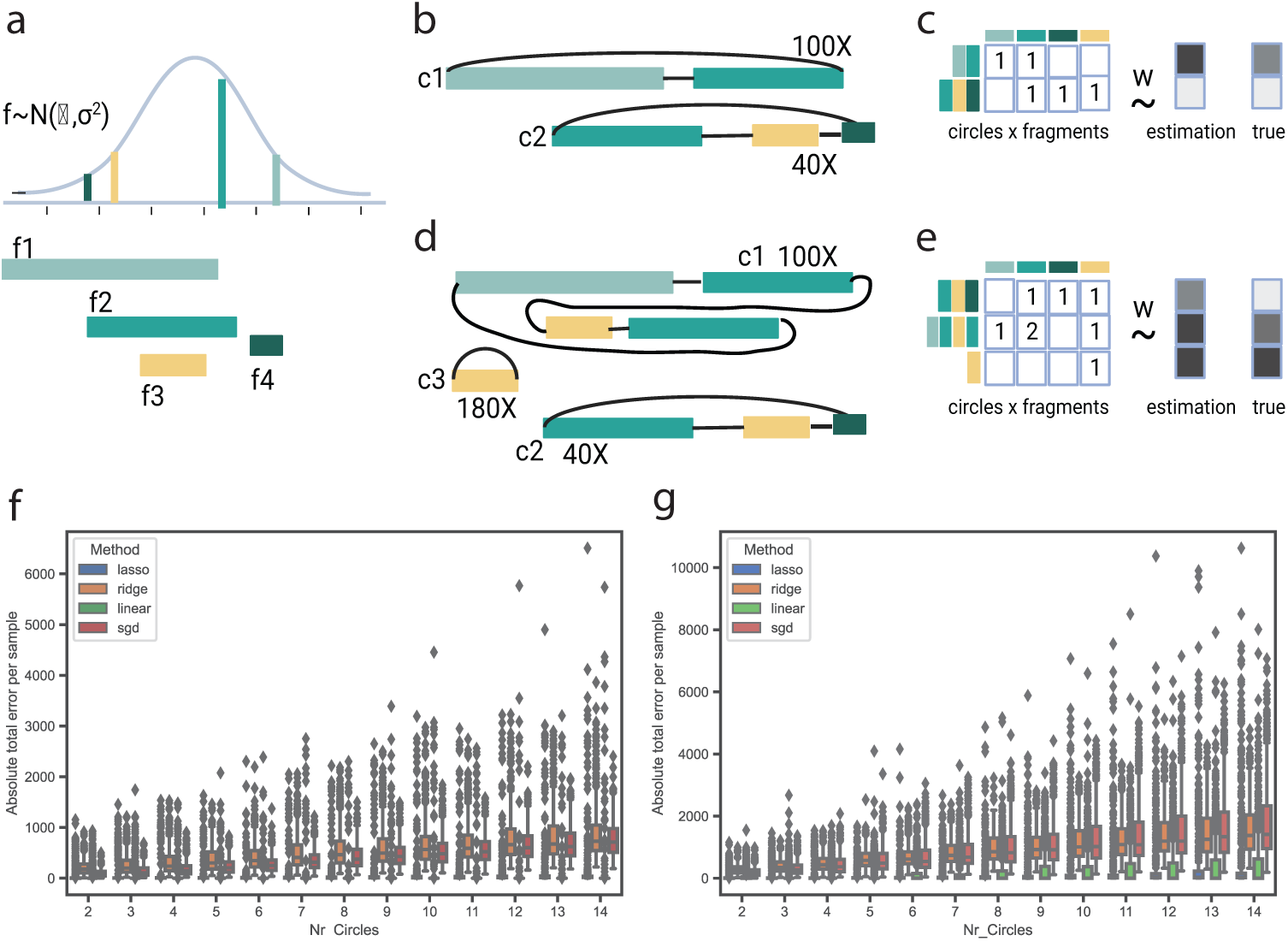
Linear models comparison to deconvolve ecDNA elements from simulated overlapping fragments data. (a) Fragments f1-4 lengths are sampled from a normal distribution N(7000,3000). (b) EcDNA elements examples (c1,c2) with overlapping fragments and amplicons copies of 100x and 40x. C1 and c2 contain unique fragments. (c,e) Schematic of the matrix formulation of the regression in (b). (d) Complex scenario of ecDNA elements with overlapping fragments. c1 contains a duplicated fragment (turquoise) and c3 is structurally a subcycle of c1 and c2. (f,g) Comparison of the four regression models, i.e. LASSO (blue), Ridge (orange), Linear regression (green) and SGD (red), X-axis represents the number of simulated ecDNA elements per sample and Y-axis quantifies the absolute total error distribution of the regression models fit. Boxplots show Q1(25%), Q2(median) and Q3(75%), interquartile range IQR = Q3 - Q1, and whiskers are 1.5 x IQR. (See Extended Methods).

**Suppl. Figure S7.**
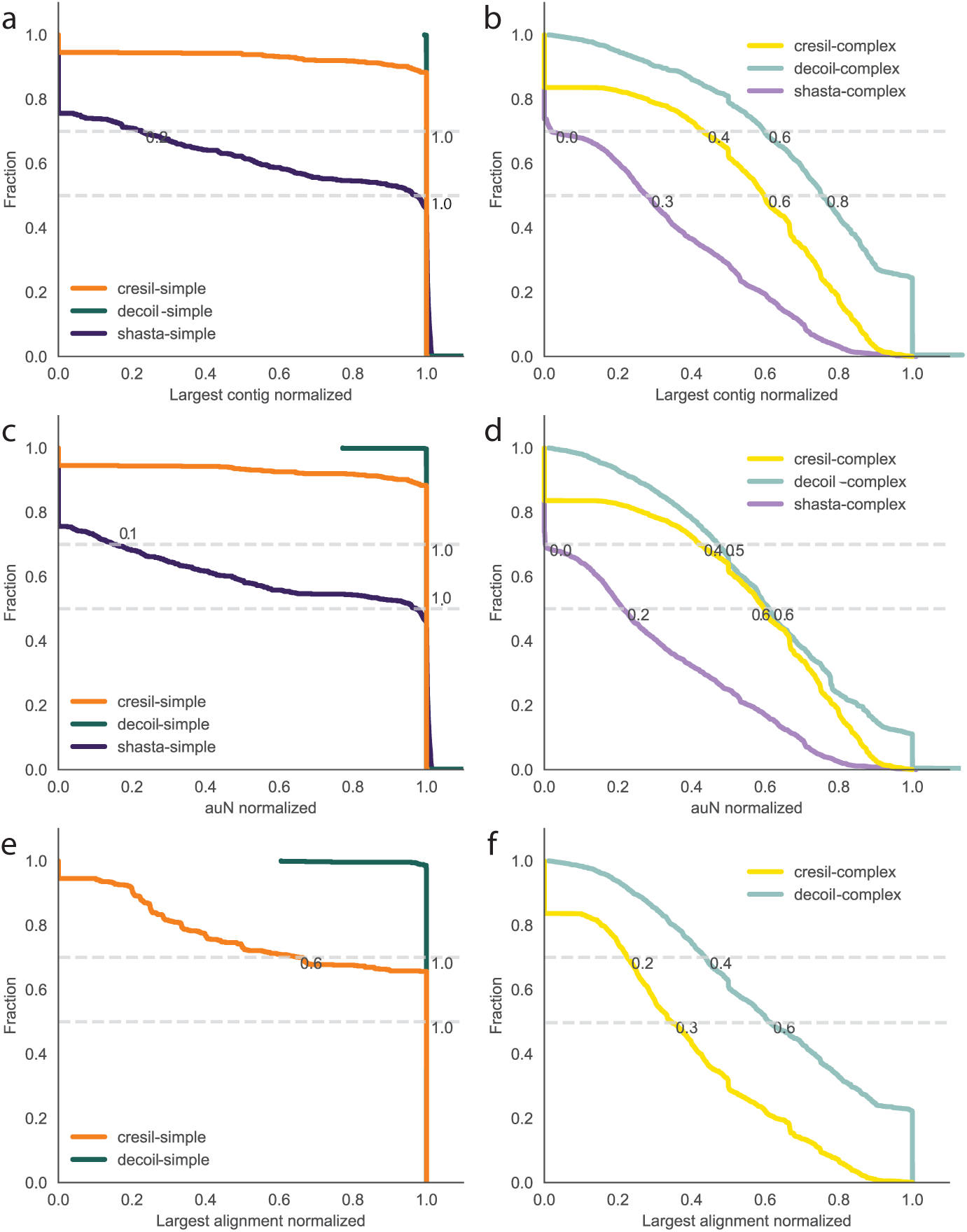
Decoil, Shasta and CReSIL comparison for the simple and complex ecDNA structures for the > 2000 simulations using different metrics. Cummulative curve of the largest contig, auN and largest alignment for (a),(c),(e) simple and (b),(d),(f) complex ecDNA simulations. (a),(b) X-axis represents the largest contig normalized by the true structure length (1 - a good reconstruction, 0 - poor reconstruction, values *>* 1 refer to reconstructions larger than the true structure). Y-axis shows the fraction of reconstructions with the specific contiguity for the three methods: Decoil (dark green / light green), CReSIL (orange / yellow), Shasta (dark purple / light purple). (c), (d) X-axis represents the auN (area under N50) normalized by the true structure length. (e), (f) X-axis represents the largest alignment normalized by the true structure length. This metric was available only for Decoil and CReSIL. The gray horizontal lines are at 0.5 and 0.7 fraction in all panels.

**Suppl. Figure S8.**
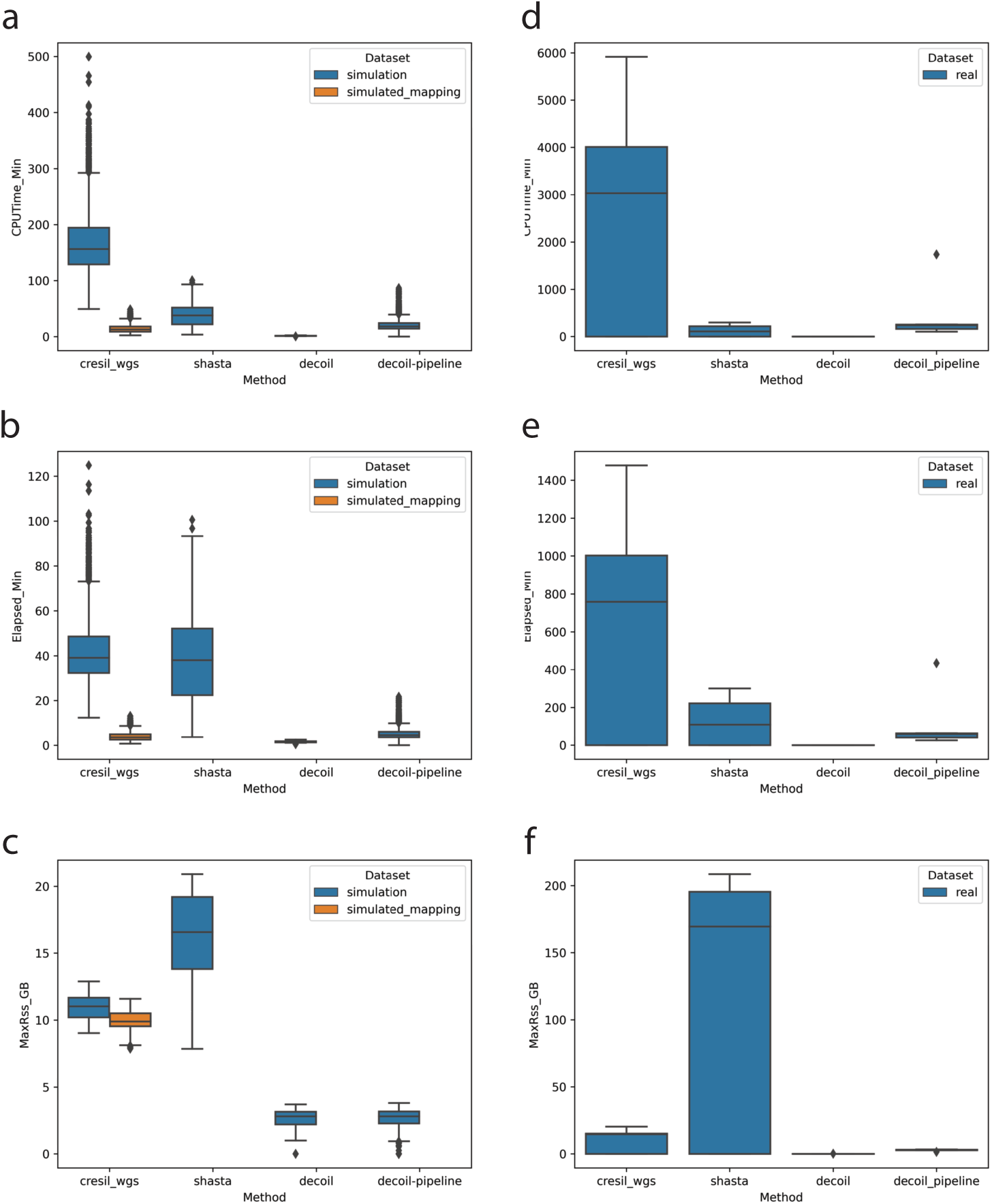
Runtime and memory benchmark. (a) CPUTime (min), (b) Elapsed time / User time (min) and (c) maximum used memory (MaxRSS) in GB for the simulated data (*>* 2000 data points). (d) CPUTime (min), (e) Elapsed time / User time (min) and (f) maximum used memory (MaxRSS) in GB for the shallow long-read whole-genome sequencing data (3-7X mean coverage, 30 data points). Shasta takes as input a .fastq. CReSIL takes as input a .fastq and performs internally alignment using minimap2 (in orange). Decoil starts with the SV calling, coverage track precomputed. Decoil-pipeline takes as input a .bam file and computes internally SV calling and the coverage track. For CReSIL, Decoil, Decoil-pipeline 4xthreads were used, for Shasta 1xthread due to intensive memory usage.

**Suppl. Figure S9.**
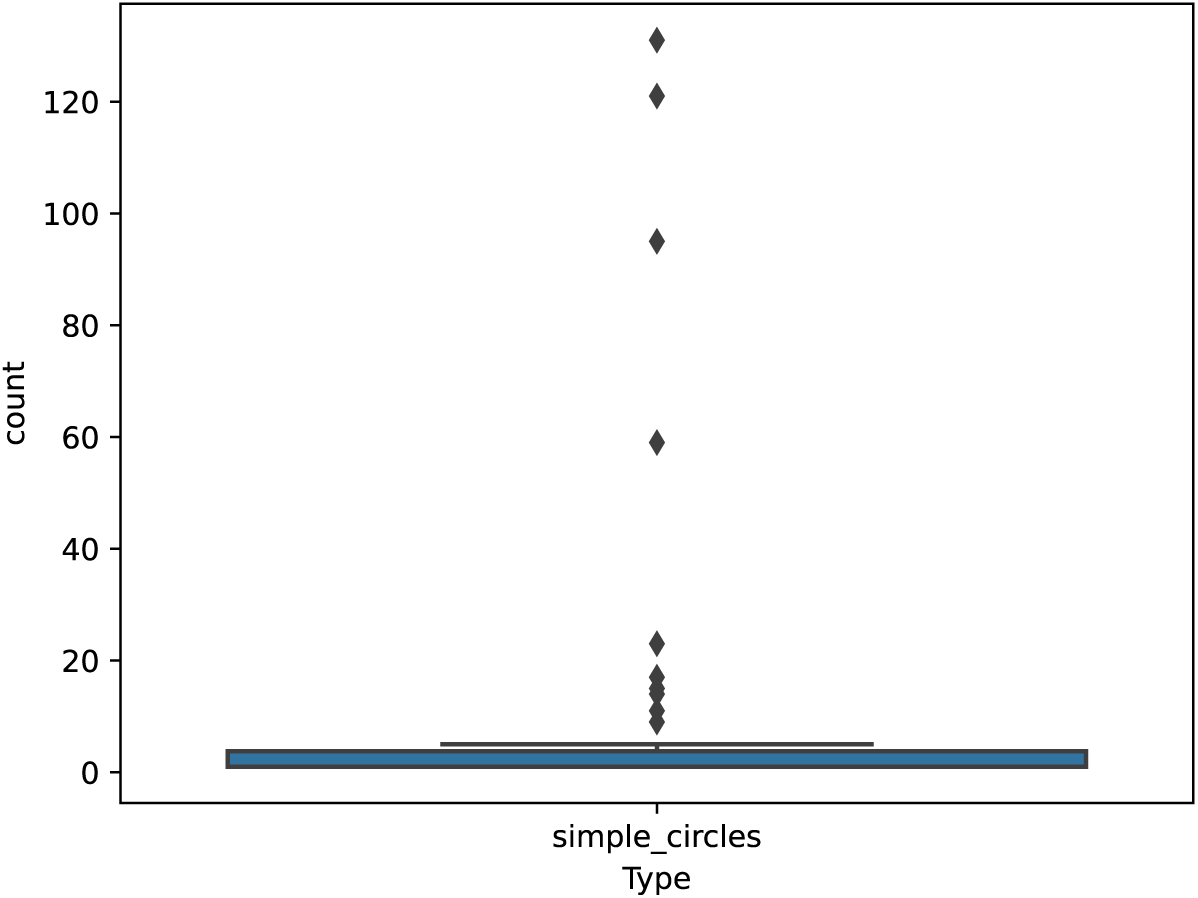
Overlapping simple cycles per cluster distribution. Y-axis (count) represents the number of simple cycles which cluster together for the real sequencing dataset, for the *MYCN* locus, i.e. have at least one overlapping fragment (mean = 8.9 and median = 1). The boxplot shows Q1(25%), Q2(median) and Q3(75%), interquartile range IQR = Q3 - Q1, and whiskers are 1.5 x IQR.

## References

[1] Kim, H. et al. Extrachromosomal DNA is associated with oncogene amplification and poor outcome across multiple cancers. Nature Genetics 52 (2020).

[2] Storlazzi, C. T. et al. MYC-containing double minutes in hematologic malignancies: Evidence in favor of the episome model and exclusion of MYC as the target gene. Human Molecular Genetics 15 (2006).

[3] Shoshani, O. et al. Chromothripsis drives the evolution of gene amplification in cancer. Nature 591 (2021).

[4] Yi, E., Chamorro González, R., Henssen, A. G. & Verhaak, R. G. Extrachromosomal DNA amplifications in cancer (2022).

[5] Pecorino, L. T., Verhaak, R. G., Henssen, A. & Mischel, P. S. Extrachromosomal DNA (ecDNA): an origin of tumor heterogeneity, genomic remodeling, and drug resistance (2022).

[6] Helmsauer, K., et al. Enhancer hijacking determines intra- and extrachromosomal circular MYCN amplicon architecture in neuroblastoma (2019).

[7] Hung, K. L. et al. Targeted profiling of human extrachromosomal DNA by CRISPR-CATCH. Nature Genetics (2022).

[8] Chamorro González, R., et al. Parallel sequencing of extrachromosomal circular DNAs and transcriptomes in single cancer cells. Nature Genetics (2023).

[9] Verhaak, R. G., Bafna, V. & Mischel, P. S. Extrachromosomal oncogene amplification in tumour pathogenesis and evolution (2019).

[10] Koche, R. P., et al. Extrachromosomal circular DNA drives oncogenic genome remodeling in neuroblastoma (2020).

[11] Prada-Luengo, I., Krogh, A., Maretty, L. & Regenberg, B. Sensitive detection of circular DNAs at single-nucleotide resolution using guided realignment of partially aligned reads. BMC Bioinformatics 20 (2019).

[12] Zhang, P., Peng, H., Llauro, C., Bucher, E. & Mirouze, M. ecc finder: A Robust and Accurate Tool for Detecting Extrachromosomal Circular DNA From Sequencing Data. Frontiers in Plant Science 12 (2021).

[13] Deshpande, V. et al. Exploring the landscape of focal amplifications in cancer using AmpliconArchitect. Nature Communications 10 (2019).

[14] Luebeck, J. et al. AmpliconReconstructor integrates NGS and optical mapping to resolve the complex structures of focal amplifications. Nature Communications 11 (2020).

[15] Wanchai, V. et al. CReSIL: accurate identification of extrachromosomal circular DNA from long-read sequences. Briefings in Bioinformatics 23 (2022).

16. Olsen, N. D. et al. precisionFDA Truth Challenge V2: Calling variants from short- and long-reads in difficult-to-map Regions. bioRxiv (2020). URL https://www.cell.com/cell-genomics/pdf/S2666-979X(22)00058-1.pdf.

[17] Olson, N. D. et al. Variant calling and benchmarking in an era of complete human genome sequences. Nature Reviews Genetics (2023).

[18] Shafin, K. et al. Nanopore sequencing and the Shasta toolkit enable efficient de novo assembly of eleven human genomes. Nature Biotechnology 38 (2020).

[19] Storlazzi, C. T. et al. Gene amplification as doubleminutes or homogeneously staining regions in solid tumors: Origin and structure. Genome Research 20 (2010).

[20] Hung, K. L. et al. ecDNA hubs drive cooperative intermolecular oncogene expression. Nature 600, 731–736 (2021).

[21] Morton, A. R. et al. Functional Enhancers Shape Extrachromosomal Oncogene Amplifications. Cell 179 (2019).

[22] Wu, S. et al. Circular ecDNA promotes accessible chromatin and high oncogene expression. Nature 575 (2019).

[23] Ono, Y., Asai, K. & Hamada, M. PBSIM2: A simulator for long-read sequencers with a novel generative model of quality scores. Bioinformatics 37 (2021).

[24] Mikheenko, A., Prjibelski, A., Saveliev, V., Antipov, D. & Gurevich, A. Versatile genome assembly evaluation with QUAST-LG, Vol. 34 (2018).

[25] Sedlazeck, F. J. et al. Accurate detection of complex structural variations using single-molecule sequencing. Nature Methods 15 (2018).

[26] Mölder, F. et al. Sustainable data analysis with Snakemake. F1000Research 10 (2021).

[27] Harase, S. On the F 2-linear relations of Mersenne Twister pseudorandom number generators. Mathematics and Computers in Simulation 100 (2014).

[28] De Coster, W., D’Hert, S., Schultz, D. T., Cruts, M. & Van Broeckhoven, C. NanoPack: Visualizing and processing long-read sequencing data. Bioinformatics 34 (2018).

[29] Ramírez, F. et al. deepTools2: a next generation web server for deep-sequencing data analysis. Nucleic Acids Research 44 (2016).

[30] Li, H. New strategies to improve minimap2 alignment accuracy. Bioinformatics 37 (2021).

