## Extended_Methods for "Decoil: Reconstructing extrachromosomal DNA structural heterogeneity from long-read sequencing data"

### Appendix B Extended methods

#### B.1 DNA extraction and nanopore sequencing

High molecular weight (HMW) DNA was extracted from 5 to 10 million cells or 15 to 25 mg of tissue using the MagAttract HMW DNA kit (Qiagen N.V., Venlo, Netherlands) according to the manufacturer's protocol. DNA concentration was measured with a Qubit 3.0 Fluorometer (Thermo Fisher) and quality control was performed using a 4200 TapeStation System (Agilent Technologies, Inc., Santa Clara, CA). For library preparation, the Ligation Sequencing Kit (SQK-LSK109 or SQK-LSK110, Oxford Nanopore Technologies Ltd, Oxford, UK) was used. All libraries were sequenced on a R9.4.1 MinION flowcell (FLO-MIN106, Oxford Nanopore Technologies Ltd, Oxford, UK) for more than 24 h.

#### Circles quantification

This steps filters quantifies the likely cycles describing the amplification. To allow reconstruction of complex structures, e.g. containing large duplications, a set of derived cycles $(D)$ was computed based on the simple cycles within a cluster. Small deletions and very rearranged genomes can lead to an increased number of high identity simple circles per cluster. A set of simple cycles $M^*$ was computed, by excluding simple cycles with an similarity higher equal than $>= x$ (default 0.9) (the longer cycle kept). The similarity was defined as the jaccard index JC:

$$JC = \frac{F1 \cap F2}{F1 \cup F2} = \frac{\sum length(fi)}{\sum length(fj)}$$

Where

- $length(fi), length(fj)$, length of fragments $fi, fj$
- Fragments $fi \in F1 \cap F2$, $fj \in F1 \cup F2$
- $F1, F2$ - fragment sets describing the cycles $c1, c2 \in M$

The derived cycles set $D$ was created by performing all the combinations between all simple cycles $c \in M^*$, which are sufficiently dissimilar in the fragment composition with $JC <= s_{max}$, (default 0.7).

To distinguish between true possible circular ecDNA elements and artifacts a *LASSO* regression is fitted against targets $Y^{|F|}$ using input $X^{|F| \times |S||D|}$, to learn the proportions of the cycles. $x_{jik} \in X$ is defined as the occurrence of fragment $f_{jk}$ in circle $c_{ik}$ and $y_{jk} \in Y$ represents the total mean coverage spanning fragment $j$, belonging to cluster $m_k$. *LASSO* was performed for each cluster $m_k$.

*LASSO* regression definition

$y_{jk} = x_{jik}\beta_{ik} + \beta_0$, $\beta_{ik}$ - weight of $c_{ik}$, $\beta_0$ - mean WGS coverage

$x_{jik} = count(f_j)$, $f_j$ occurs in cycle $c_i$,

$i = \overline{1, \mid S \mid\mid D \mid}, j = \overline{1, \mid F \mid}, k = \overline{1, \mid M \mid}$

After $LASSO$ regression fit, circles $c_{ik}$ with weight $\beta_{ik} > t$ where kept, where threshold $t$ is defined as $t = max(min(coverage(f_j))/5, 10)$. The obtained $LASSO$ coefficient $\beta_{ik}$ represent the estimated proportions of each cycle $c_{ik}$. The higher the value the more likely is the cycle to be a true ecDNA element. To avoid overfitting of the model, a penalty term $alpha = 0.1$ was used.

##### Candidates selection

From the candidates list we filter out cycles with an estimated proportions <= mean WGS coverage (default). Lastly, circular elements larger than 0.1 MB (threshold published by [13]) are labeled as ecDNA.

##### Output and Visualization

The algorithm outputs the candidates list as *.bed, *.fasta, including the mean coverage and orientation per fragment, estimated proportions of circular element. The *summary.txt* displays all found circular elements, which includes small circles and ecDNA. The reconstructions labeled as ecDNA are visualised using gGnome (https://github.com/mskilab/gGnome).

#### B.3 Ranking system of ecDNA topologies

To assess Decoil's reconstruction performance, we generated an *in-silico* collection of ecDNA elements, spanning various sequence complexities for systematic evaluation. We introduced a ranking system and defined seven topologies of increasing computational complexity, based on the SV's contained on the ecDNA element:

1. *Simple circularization* - no structural variants on the ecDNA
2. *Simple SV's* - ecDNA element contains either a series of inversions or deletions
3. *Mixed SV's* - ecDNA element has a combination of inversions and deletions
4. *Multi-region* - ecDNA element contains different genomic regions from the same chromosome (DEL, INV and TRA allowed)
5. *Multi-chromosomal* - ecDNA element originates from multiple chromosomes (DEL, INV and TRA allowed)
6. *Duplications* - ecDNA element contains duplications defined as a region larger than 50 bp repeated on the amplicon (DUP's + other simple rearrangements)
7. *Foldbacks* - ecDNA element contains a foldback defined as a two consecutive fragments which overlap in the genomic space, with different orientations (INVDUP's + all other simple SV's)

Every topology can contain a mixture of all other low-rank topologies.

#### B.4 Simulate ecDNA sequence templates

The simulation framework contains probabilistic variables, which model the chromosome weights, fragment position, fragment length, small deletion ratio, inversion ratio, foldback ratio, and tandem-duplication ratio. Simulation strategy for individual ecDNA templates starts by choosing the genomic position relative to previous simulated fragment, covering four scenarios (Figure 2a#1):

- neighbor - the next simulated fragment starts right next to the previous fragment
- [0 to 5 kb] - the next simulated fragment starts within a 5 kb distance relative to the previous fragment
- >5 kb - the next simulated fragment starts a least at a 5 kb distance relative to the previous fragment
- switch chromosome - the next simulated fragment is sampled from another chromosome

Next, to simulate small deletions (DELs), $< 10\%$ of fragment size can be cut out with a certain probability p, at the left or right end of the fragment (Figure 2a#2). With a probability p, inversions (Figure 2a#3) and tandem-duplications (Figure 2a#4) are simulated. To cover a wide range of possible conformations we generate first a so-called conformation array, which encodes the different event types for describing the simulation of individual ecDNA template. The conformation array has binary entries (except the first position which encodes the fragments number), where every bit is set to 0 (disable the occurrence of the event on ecDNA) or to 1 (allows the occurrence of the event with a probability p).

**Supplementary Table S5** Conformation array for ecDNA templates simulation. N_FRAG - number of fragments; SMALL_DEL - allow small deletions on the right and left side of the fragment; DUP - allow simple duplication; INV - allow inversions; INTERCHR - allow fragments to originate from multiple chromosomes; MULTI_REGION - allow fragments to originate from multiple regions on same chromosome; FOLDBACK - allow foldbacks, which are here defined as two overlapping genomic fragments, which are immediately chained in the ecDNA template, regardless of the strand orientation.

| N_FRAG | SMALL_DEL | DUP | INV | INTERCHR | MULTI_REGION | FOLDBACK |
|---|---|---|---|---|---|---|

Conformation array for the seven topologies based on which multiple rounds of simulations were performed:

Simple circularization:

| | | | | | | |
|---|---|---|---|---|---|---|
| 1 | 0 | 0 | 0 | 0 | 0 | 0 |

Simple SV's:

| | | | | | | |
|---|---|---|---|---|---|---|
| 2 - 10 | [0\|1] | 0 | 0 | 0 | 0 | 0 |

Mixed SV's:

| | | | | | | |
|---|---|---|---|---|---|---|
| 2 - 10 | 1 | 0 | 1 | 0 | 0 | 0 |

Multi-region :

| | | | | | | |
|---|---|---|---|---|---|---|
| 2 - 10 | [1\|0] | 0 | [1\|0] | 0 | 1 | 0 |

Multi-chromosomal:

| | | | | | | |
|---|---|---|---|---|---|---|
| 2- 10 | [1\|0] | 0 | [1\|0] | 1 | 1 | 0 |

Duplications:

| | | | | | | |
|---|---|---|---|---|---|---|
| 2 - 10 | [1\|0] | 1 | [1\|0] | [1\|0] | [1\|0] | 0 |

Foldbacks:

| | | | | | | |
|---|---|---|---|---|---|---|
| 2 - 10 | [1\|0] | [1\|0] | [1\|0] | [1\|0] | [1\|0] | 1 |

In total 577 conformation arrays were obtained, based on which more than 2000 ecDNA templates were generated. Code available under https://github.com/madagiurgiu25/ecDNA-sim.

#### B.5 Simulate *in-silico* long-read ecDNA-containing samples

To assess ecDNA reconstruction performance, *in-silico* ecDNA-containing samples were generated based on the ecDNA sequence templates collection. The workflow takes as input the defined ecDNA elements in .bed format and generates its associated .fasta reference. Afterwards, noisy long-reads, with an average length of 7,000 bp, are sampled from this reference using an adapted version of PBSIM2 (Ono et al. 2021 [23]), at a specified depths of coverage. This package was customized for the purpose of this paper to (1) allow reads sampling from a circular reference, and (2) provide a better coverage uniformity of the reads at fragments boundary by using Mersenne twister (Harase 2014 [27]) instead of the pseudorandom number generator included in the original package (https://github.com/madagiurgiu25/pbsim2). The *in-silico* reads are stored in .fastq format. This workflow steps is part of the benchmarking pipeline https://github.com/madagiurgiu25/ecDNA-simulate-validate-pipeline.

#### B.6 Alignment-free ecDNA reconstruction using Shasta from simulated data

To *de-novo* assemble the simulated ecDNA the reads were filtered using NanoFilt [28] 2.6.0 (-l 300 -q 20 –headcrop 20 –tailcrop 20). *De-novo* assembly was performed using Shasta [18] 0.10.0 with parameters –config Nanopore-May2022 –Reads.minReadLength 1000 –Kmers.distanceThreshold 500 –Kmers.probability 0.5.

#### B.7 Alignment-based ecDNA reconstruction using Decoil from simulated data

To reconstruct the ecDNA using Decoil, the reads were filtered using NanoFilt [28] 2.6.0 (-l 300 -q 20 –headcrop 20 –tailcrop 20), aligned to the reference genome GRCh38/hg38 using ngmlr [25] 0.2.7 with standard parameters. Structural variant calling was performed using sniffles [25] 1.0.12 (–min_homo_af 0.7 –min_het_af 0.1 –min_length 50 –cluster –min_support 4) and the bigWig coverage tracks were computed using bamCoverage (-50 bins) from deepTools [29] 3.5.1 suite. Decoil used the alignment, SV calls and coverage profile as input to reconstruct simulated ecDNA (–min-vaf 0.01 –min-cov-alt 6 –min-cov 8 –max-explog-threshold 0.01 –fragment-min-cov 10 –fragment-min-size 500).

#### B.8 Alignment-based ecDNA reconstruction using CReSIL from simulated data

Simulated ecDNA was also identified using CReSIL [15] v1.0.0 [https://github.com/visanuwan/cresil, commit:646aec9], with standard parameters, using 'cresil trim', followed by 'cresil identify_wgls' and reference genome GRCh38/hg38.

#### B.9 Linear models comparison to deconvolve ecDNA elements from simulated overlapping fragments data

In order to identify the probable ecDNA components within a given sample, we employ a regression-based technique to deconvolve the circular paths that best align with the coverage profile and determine their estimate proportions. Four different regression models were tested, i.e LASSO (Least Absolute Shrinkage and Selection Operator), Rigde regression, Linear regression and SGD (Stochastic Gradient Descent) regression. This experiment simulates the matrix formulation for the regression. In the simulation it can be specified how many fragments two different circular structure share and randomly chosen. The amplicon copies per circular structure is sampled from a normal distribution $\mathcal{N}(200, 150)$. For every fragment of the circular the length is samples from a normal distribution $\mathcal{N}(7000, 3000)$. To calculate the error between predicted coverage $Y_p$ and true coverage profile $Y_t$ the total absolute error was used, i.e. $e = \sum_i |y_{pi} - y_{ti}|$, with $y_{pi} \in Y_p$ and $y_{ti} \in Y_t$.

#### B.10 Performance evaluation on simulated data

To evaluate the correctness of reconstruction for both, Decoil (alignment-based) and Shasta (alingment-free), Quast [24] 5.2.0 was applied to compute different metrics (https://quast.sourceforge.net/docs/manual.html). The overall reconstruction performance was quantified as the mean and standard deviation of the largest contig metric, defined as the longest contig in the assembly. The contiguity of the reconstruction was visualized using dotplots, for which paf alignments from the true and reconstructed were generated (for both Decoil and Shasta) using minimap2 [30] 2.26-r1175. Definition of the metrics which is an adaptation of Quast definitions.:

- Largest_contig_norm - Largest contig is the length of the longest contig in the assembly, normalized by true length
- Total_length_norm - is the total number of bases in the assembly, normalized by true length
- N50_norm - is the length for which the collection of all contigs of that length or longer covers at least half an assembly, normalized by true length
- N90_norm - same as N50 but but with 90% instead of 50%
- auN_norm - is the area under the N50, normalized by true length. This metric was proposed and justified by Heng Li in his blog https://lh3.github.io/2020/04/08/a-new-metric-on-assembly-contiguity.
- Largest_alignment_norm - is the length of the largest continuous alignment in the assembly, normalized by true length

- Total_aligned_length_norm - is the total number of aligned bases in the assembly, normalized by true length
- Misassembled_contigs_length_norm - is the total number of bases in misassembled contigs, normalized by true length

#### B.11 Evaluate amplicon's breakpoints recovery in ecDNA mixtures

To evaluate how well Decoil can reconstruct ecDNA elements with overlapping footprints a series of dilutions was generated by mixing the CHP212, STA-NB-10DM and TR14 cell lines at different ratios. We generated two types of mixtures. First, 100% of one sample with different percentages of another sample, i.e. 10, 25, 50, 75, 90, 100% (Figure 3c) were combined. Secondly, mixtures at different ratios for both samples (10-90, 25-75, 50-50, 75-25, 90-10%) were generated. Picard 2.26 (https://broadinstitute.github.io/picard/) was used to downsample the .bam files to 10, 25, 50, 75, 90% and samtools 1.9 to merge the different ratios and to create *in-silico* ecDNA mixtures. SV calling was performed using sniffles [25] 1.0.12 with same parameters as for the original 100% .bam files, i.e. –min_homo_af 0.7 –min_het_af 0.1 –min_length 50 –cluster –min_support 4. Decoil was run on all these mixtures with parameters –min-vaf 0.01 –min-cov-alt 10 –min-cov 10 –max-explog-threshold 0.01 –fragment-min-cov 10 –fragment-min-size 500. The completeness of the reconstructed ecDNA elements in mixtures was evaluated by counting how many breakpoints are identical compared to the true ecDNA elements in the 100% samples.

#### B.12 Preprocess nanopore sequencing data from cell lines and patient samples

For the analysis five neuroblastoma cell lines and 13 patients were sequenced using shallow whole-genome sequencing. For all the samples the status of MYCN amplification on ecDNA was experimentally determined by FISH. The patients cohort included 10 patients ecDNA positive and three ecDNA negative, serving as negative control. One ecDNA-containing sample was removed from the analysis due to failed QC. The cell lines CHP212, TR14, STA-NB-10DM and all 13 patient samples were preprocessed by performing base-calling using Guppy 5.0.14 (dna_r9.4.1_450bps_hac model), followed by a quality check using NanoPlot 1.38.1. The reads were filtered by quality using NanoFilt [28] 2.8.0 (-l 300 –headcrop 50 –tailcrop 50) and aligned using ngmlr [25] 0.2.7 against the reference genome GRCh38/hg38. The structural variant calling was performed using sniffles [25] 1.0.12 (–min_homo_af 0.7 –min_het_af 0.1 –min_length 50 –min_support 4). The bigWig coverage tracks were obtained by applying bamCoverage (-50 bins) from deepTools [29] 3.5.1 suite. The cell lines LAN-5 and CHP126 were similarly processed using the reference genome GRCh37/hg19. The pipeline is available under https://github.com/henssen-lab/nano-wgs.

#### B.13 Reconstruct ecDNA elements for cell lines and patient samples using Decoil

To reconstruct the ecDNA elements for CHP212, TR14 and STA-NB-10DM Decoil was applied using the parameters –min-vaf 0.1 –min-cov-alt 10 –min-cov 8 –fragment-min-cov 10 –fragment-min-size 1000 –filter-score 35 or –min-vaf 0.01 –min-cov-alt 10 –min-cov 10 –max-explog-threshold 0.01 –fragment-min-cov 10 –fragment-min-size 500, the reference genome GRCh38/hg38 and annotation GENCODE v42. Similarly, for LAN-5 and CHP126 the ecDNA reconstruction was performed using Decoil with same parameters, reference genome GRCh19/hg19 and annotation GENCODE v41. The ecDNA elements in patient samples were reconstructed by Decoil using –min-vaf 0.1 –min-cov-alt 10 –min-cov 30 –max-explog-threshold 0.01 –fragment-min-cov 20 –fragment-min-size 100.

#### B.14 Patient sample and clinical access

Patients were registered and treated according to the trial protocols of the German Society of Pediatric Oncology and Hematology (GPOH). This study was conducted in accordance with the World Medical Association Declaration of Helsinki (2013) and good clinical practice; informed consent was obtained from all patients or their guardians. The collection and use of patient specimens was approved by the institutional review boards of Charité-Universitätsmedizin Berlin and the Medical Faculty, University of Cologne. Specimens and clinical data were archived and made available by Charité-Universitätsmedizin Berlin or the National Neuroblastoma Biobank and Neuroblastoma Trial Registry (University Children's Hospital Cologne) of the GPOH. The *MYCN* gene copy number was determined as a routine diagnostic method using FISH.

#### B.15 Runtime and memory benchmarking

For both simulated and real datasets, we conducted an analysis of the runtime and memory usage. The runtime, including the raw elapsed time (ElapsedRaw) and CPU time (CPUTime), was measured. Additionally, memory usage was assessed using the maximum resident set size (MaxRss). These metrics were derived from the Slurm output, providing insights into the computational resources consumed during the analysis.
